# Differentially accessible Cdc4 phospho-degrons regulate Ctf19^CCAN^ kinetochore subunit stability in mitosis

**DOI:** 10.1101/2021.02.12.430993

**Authors:** Miriam Böhm, Kerstin Killinger, Alexander Dudziak, Pradeep Pant, Karolin Jänen, Simone Hohoff, Karl Mechtler, Mihkel Örd, Mart Loog, Elsa Sanchez-Garcia, Stefan Westermann

## Abstract

Kinetochores are multi-subunit protein assemblies that link chromosomes to microtubules of the mitotic and meiotic spindle. How effective, yet strictly centromere-dependent kinetochore assembly is coupled to cell cycle progression is incompletely understood. Here, by combining comprehensive phosphorylation analysis of native Ctf19^CCAN^ subunits with biochemical and functional assays in the model system budding yeast, we demonstrate that Cdk1 phosphorylation activates phospho-degrons on the essential subunit Ame1^CENP-U^ which are recognized by the E3 ubiquitin ligase complex SCF-Cdc4. Gradual phosphorylation of degron motifs culminates in M-Phase and targets the protein for degradation. Binding of the Mtw1 complex shields the proximal phospho-degron, protecting kinetochore-bound Ame1 from the degradation machinery. Artificially increasing degron strength partially suppresses the temperature-sensitivity of a *cdc4* mutant, while overexpression of Ame1-Okp1 is toxic to cells, demonstrating the physiological importance of this mechanism. We propose that phospho-regulated clearance of excess CCAN subunits protects against ectopic kinetochore assembly and contributes to mitotic checkpoint silencing. Our results suggest a novel strategy for how phospho-degrons can be used to regulate the assembly of multi-subunit complexes.

## Introduction

Kinetochores form a dynamically regulated binding interface between chromosomes and the mitotic spindle, a connection which is essential to partition the genetic material equally during cell division (Biggins, 2013). As chromatin-bound multi-protein complexes, kinetochores assemble exclusively at centromeres. Biochemical experiments have elucidated the connectivity between multiple subcomplexes in the context of an assembled kinetochore (Musacchio and Desai, 2017), and structural biology has started to define the organization of these building blocks at atomic resolution (Hinshaw and Harrison, 2019; Yan et al., 2019). It is, however, still poorly understood how efficient - yet strictly centromere-dependent - kinetochore assembly is accomplished from hundreds of individual building blocks in a cell cycle dependent manner.

In budding yeast centromeres are attached to kinetochore microtubules during almost the entire vegetative cell cycle. Upon replication, kinetochores assemble quickly on both sister chromatids to ensure effective bi-orientation. Live cell imaging experiments suggest that kinetochore assembly upon replication of centromere DNA occurs rapidly and is completed within 10-15 min (Tanaka et al., 2007). This is remarkable as even the relatively simple kinetochores in budding yeast consist of more than 40 different proteins. These protein subunits assemble in two layers: an inner centromere-binding layer, whose subunits contact the centromeric nucleosome and are collectively called the Constitutive Centromere Associated Network (CCAN, or Ctf19^CCAN^ in budding yeast) and a microtubule-binding layer containing the KMN network. The CCAN consists of the essential homodimeric protein Mif2^CENP-C^, which contacts and organizes many other subunits of the CCAN (Klare et al., 2015) and also of the COMA complex, which consists of the conserved subunits Ctf19^CENP-P^, Okp1^CENP-Q^, Mcm21^CENP-O^ and Ame1^CENP-U^, of which Ame1 and Okp1 are essential for cell viability in budding yeast (Hornung et al., 2014). The CCAN directly binds to a specialized nucleosome by recognizing the H3 variant Cse4 via the Mif2 and Ame1-Okp1 subunits (Anedchenko et al., 2019; Fischbock-Halwachs et al., 2019; Killinger et al., 2020). The KMN network, on the other hand is composed of Knl1 (Spc105 in yeast), Mis12c (Mtw1c in yeast) and Ndc80c (Ndc80c in yeast). These three subcomplexes are necessary to efficiently couple dynamic microtubules to centromeres (Cheeseman et al., 2006) and also form a regulated recruitment platform for components of the Spindle Assembly Checkpoint (SAC) (London et al., 2012). The outer yeast kinetochore also includes a ring-forming protein complex called the Dam1c, which is required for robust coupling of the KMN network to dynamic microtubules (Lampert et al., 2013; Tien et al., 2010).

During kinetochore formation, cells have to strike a balance between allowing efficient assembly at the centromere, while at the same time preventing the formation of kinetochores at non-centromeric loci. This is best exemplified by the histone variant CENP-A. High levels of CENP-A facilitate its mis-incorporation into non-centromeric loci (Heun et al., 2006; Van Hooser et al., 2001) and can promote the formation of ectopic kinetochores and genetic instability (Amato et al., 2009). Likewise, artificial recruitment of CCAN subunits like CENP-C or CENP-T to non-centromeric loci can promote the formation of ectopic kinetochores (Gascoigne et al., 2011). In budding yeast, Cse4^CENP-A^ protein levels are controlled via regulated proteolysis involving the E3 ubiquitin ligase Psh1, which prevents mis-incorporation of Cse4 into chromosome arms (Ranjitkar et al., 2010). Whether such regulatory systems operate for CCAN subunits is currently not known. Moreover, there is evidence that dynamic changes in the architecture of the budding yeast kinetochores occur, with subunits being added during anaphase (Dhatchinamoorthy et al., 2017; Dhatchinamoorthy et al., 2019). How these dynamic changes are regulated to assemble kinetochores with the correct subunit stoichiometry, while avoiding ectopic kinetochore assembly at the same time, is not well understood.

Phospho-regulation plays an important role for multiple aspects of kinetochore function. The Ipl1^AuroraB^ kinase regulates kinetochore-microtubule attachments during error correction to ensure sister chromatid bi-orientation and it also promotes kinetochore assembly by phosphorylating the Mis12c subunit Dsn1 (Dimitrova et al., 2016). The major regulator of the mitotic cell cycle Cdc28^Cdk1^ on the other hand, promotes outer kinetochore assembly in human cells the by stimulating the phospho-dependent recruitment of Ndc80 via CENP-T (Huis In ’t Veld et al., 2016; Nishino et al., 2013). By contrast, in budding yeast the Cnn1-Ndc80c interaction is negatively regulated by Mps1 phosphorylation (Malvezzi et al., 2013). Apart from these examples, however, it is still unclear, how phospho-regulation is linked to cell cycle progression and which aspects of kinetochore function may be affected.

Here we set out to define mechanisms of phospho-regulation in the context of the budding yeast inner kinetochore. We identify multiple Cdk1 substrates in the yeast Ctf19^CCAN^ complex and demonstrate that a subset of these sites constitute phospho-degron motifs that are activated in a cell cycle dependent manner and recognized by the SCF-Cdc4 E3 ubiquitin ligase complex. Our results reveal a molecular link between the core cell cycle machinery and key protein subunits of the inner kinetochore that serves to couple kinetochore subunit turnover to cell cycle progression.

## Results

### Mapping of native phosphorylation sites identifies multiple candidate Cdk1 substrates in the yeast CCAN

To investigate the phosphorylation status of native yeast CCAN subunits, we affinity-purified TAP-tagged components representing major CCAN subcomplexes (Chl4^CENP-N^-TAP, Mcm16^CENP-H^-TAP, Cnn1^CENP-T^-TAP, Mhf1^CENP-S^-TAP, Mhf2^CENP-X^-TAP) from log-phase yeast extracts and identified phosphorylation sites by mass spectrometry. In total, this analysis detected more than 70 phosphorylation sites on nine different CCAN subunits (**Table 1, Supplementary Data File 1**). Of these sites, nine followed the minimal consensus motif for Cdk1 phosphorylation (S/TP), while four sites (in the subunits Mcm21, Nkp1 and Cnn1) followed the full Cdk1 consensus (S/TP_K/R). While Cdk1 phosphorylation of Cnn1 has been described before (Bock et al., 2012; Schleiffer et al., 2012), our analysis identified Ame1 as a candidate Cdk1 target, with a cluster of four minimal Cdk1 sites being phosphorylated in the N-terminus of the protein.

**Table 1:** Analysis of Ctf19c^CCAN^ phosphorylation in yeast extracts. Native CCAN phosphorylation sites detected after purification of TAP-tagged kinetochore subunits from yeast extracts. For details see Supplementary Data File 1.

| S.c.CCAN subunit | Human homolog | % sequence coverage | Total P-sites detected | Minimal Cdk1 sites detected (S/T)P | Full Cdk1 sites detected (S/TP_K/R) |
| --- | --- | --- | --- | --- | --- |
| Ame1 | CENP-U | 79 | 8 | 4 (T31, S41, S45, S53) | - |
| Okp1 | CENP-Q | 85 | 6 | - | - |
| Mcm21 | CENP-O | 92 | 3 | - | 1 (S139) |
| Ctf19 | CENP-P | 80 | 1 | - | - |
| Nkp1 | - | 97 | 3 | - | 1 (S222) |
| Nkp2 | - | 89 | - | - | - |
| Chl4 | CENP-N | 98 | 3 | 1 (S281) | - |
| Iml3 | CENP-L | 95 | - | - | - |
| Ctf3 | CENP-I | 75 | - | - | - |
| Mcm22 | CENP-H | 97 | - | - | - |
| Mcm16 | CENP-K | 93 | - | - | - |
| Cnn1 | CENP-T | 90 | 17 | 2 (T42, S192) | 2 (T21, S177) |
| Mhf1 | CENP-S | 93 | 3 | 1 (T34) | - |
| Mhf2 | CENP-X | 95 | 1 | 1 (S60) | - |

### The essential CCAN subunit Ame1^CENP-U^ is a Cdk1 substrate in vivo and in vitro

The cluster of Cdk1 phosphorylation sites in Ame1 is located close to the functionally important Mtw1c binding domain (**Figure 1A**). We phosphorylated recombinant Ame1-Okp1 complex with purified Cdc28-Clb2 (M-Cdk1) or Cdc28-Clb5 (S-Cdk1) in vitro and mapped phosphorylation sites by mass spectrometry (**Supplementary Data File 2**). Ame1 displayed a noticeable shift in migration in SDS-PAGE upon phosphorylation with Cdc28-Clb2, but less so with Cdc28-Clb5 (**Figure 1B**). Quantitative phosphorylation analysis confirmed that Ame1 is a preferred M-Cdk1 substrate, whereas Okp1 is a better substrate for the S-Cdk1 complex Cdk1-Clb5. The phosphorylation of Okp1 was promoted by the hydrophobic patch, a known substrate docking region in Clb5 and Clb2 cyclins (**Supplementary Figure 1**). The mapped phosphorylation sites closely corresponded to the sites detected on native Ame1, in particular phosphorylation of the residues Thr31, Ser41, Ser45, and Ser52/Ser53, was both detected in vivo and in vitro (**Figure 1C**). In case of Ser52/Ser53 either one, or both adjacent sites may be phosphorylated. For the subsequent analysis, we focused on seven N-terminal phosphorylation sites, as the two C-terminal sites Ser277 and Ser323 were not phosphorylated by Cdc28-Clb2 in vitro. To analyze the functional role of Ame1 phosphorylation we mutated the cluster to either alanine (Ame1-7A) or glutamic acid (Ame1-7E) to eliminate or mimic phosphorylation, respectively. We integrated Flag-tagged Ame1 constructs under their endogenous promoter with these mutations into yeast and analyzed cell extracts by western blotting. Analysis of log phase extracts showed that wild-type Ame1 displayed multiple slowly migrating forms which were eliminated in the 7A mutant (**Figure 1D**). By contrast, Ame1-7E migrated much more slowly than wild-type, its position in SDS-PAGE corresponding to the most slowly migrating forms of Ame1-WT. Ame1-7A and -7E mutants were viable when expressed as the sole source of Ame1 in the cell. In an anchor-away approach, in which endogenous Ame1 is removed from the nucleus upon addition of rapamycin, both Ame1-7A and -7E variants supported viability with little difference in growth rate on rich media compared to wild-type Ame1 (**Figure 1E**). Analysis of internal truncations, which maintained the essential Mtw1c-binding N-terminus (residues 1-15) showed that deleting the region harboring the entire phospho-cluster (Δ31-116) yielded a slow growth phenotype, while a more extensive deletion was inviable (**Figure 1F**). We conclude that Ame1 phosphorylation is not required for viability, but the N-terminus contributes to an important aspect of Ame1 function, even when the Mtw1-binding domain is retained.

**Figure 1:**
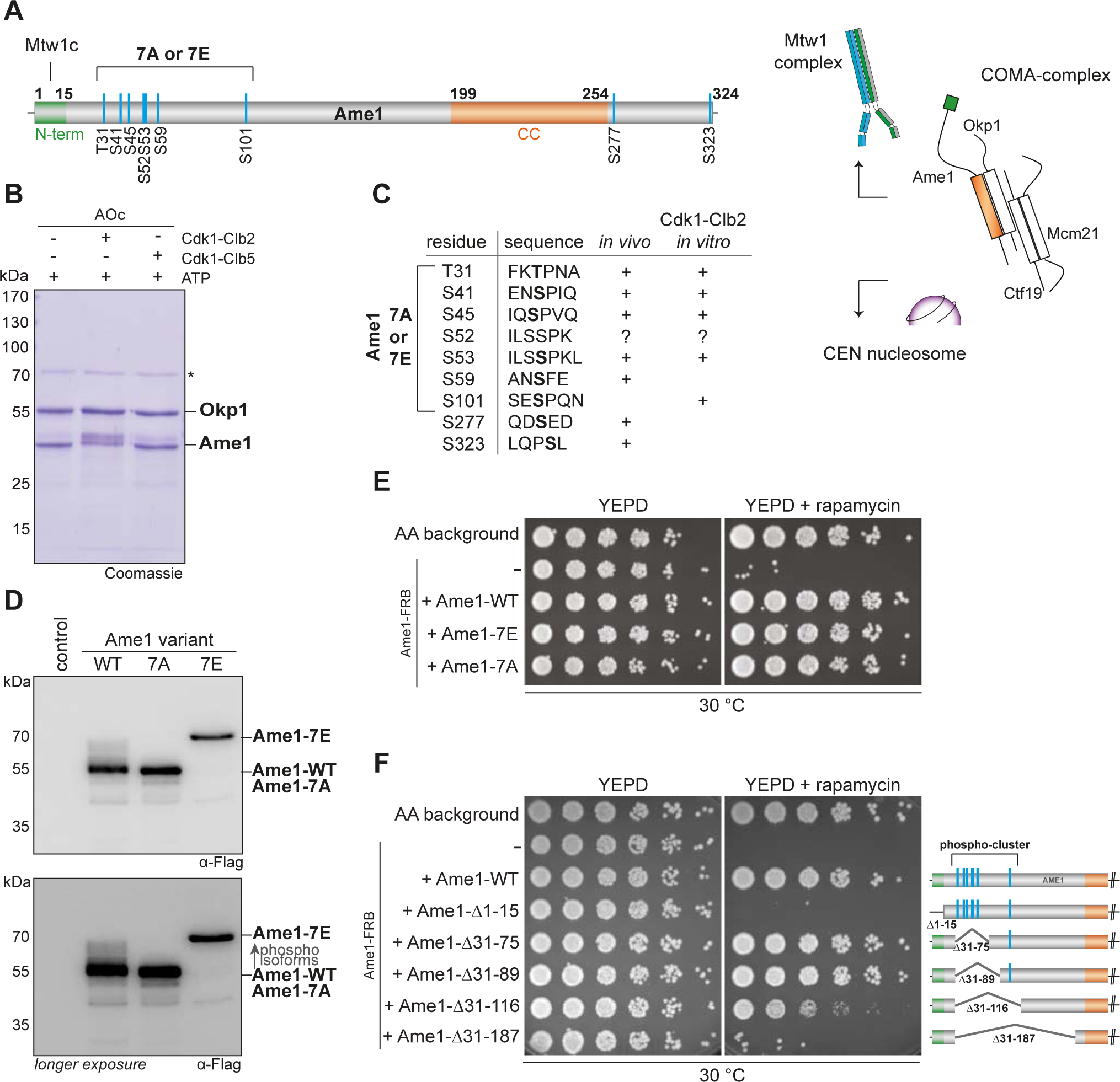
Phosphorylation analysis of the essential CCAN subunit Ame1. **A:** Organization of the essential CCAN component Ame1^CENP-U^ and localization of phosphorylation sites. Ame1 shows a Cdk1 phosphorylation cluster (T31, S41, S45, S52, S53, S59, S101) at the N-terminus. The first 15 amino acids are essential for Mtw1c binding, the coiled-coil region (aa 199-254) is required for heterodimerization with Okp1^CENP-Q^. **B:** In vitro kinase assay with recombinant Ame1-Okp1c with either Cdk1-Clb2 or Cdk1-Clb5. The migration pattern of Ame1 is shifted to a slowly migrating form when incubated with Cdk1-Clb2. **C:** List of all mapped Ame1 phosphorylation sites either in vivo or in vitro. T31, S41, S45, S53, S101 show the minimal motif for Cdk1 (S/TP). **D:** Stably integrated Ame1-variants display distinct migration patterns in SDS-PAGE. Ame1-WT shows multiple slowly migrating forms that are eliminated in Ame1-7A and Ame1-7E. **E:** Serial dilution assay of Ame1-variants using the FRB anchor-away system. Ame1-WT and both mutants can rescue the growth defect, when endogenous Ame1 is anchored away from the nucleus. **F:** Serial dilution assay of internal Ame1 truncation mutants in the anchor-away system.

### Non-phosphorylatable Ame1 mutants accumulate to increased protein levels

During our cellular characterization experiments for Ame1 phospho-mutants we expressed wild-type or mutant versions of Ame1, along with its binding partner Okp1 from a two-micron plasmid under control of a Galactose-inducible promoter (**Figure 2A**). Western Blot analysis showed that wild-type Ame1 gradually accumulated over the course of five hours after switching the cells to Galactose. Strikingly, the non-phosphorylatable Ame1-7A mutant accumulated to much higher protein levels in the same time span, leading to a roughly four-fold increase in steady state level compared to wild-type after five hours in Galactose (**Figure 2B and C**). In this experiment, the Ame1-7E mutant behaved similar to the -7A mutant, suggesting that it may constitute a phospho-preventing rather than a phospho-mimetic mutation (**Figure 2B and C**). By contrast, Okp1 expressed from the same plasmid showed no change in protein level in the different Ame1 mutants, arguing that differences in plasmid stability or mitotic retention cannot be the cause for the observed effect on the Ame1 protein level. Protein level differences between Ame1 phospho-mutants were also observed when overexpression was performed in a *mad1Δ* strain background (**Supplementary Figure 2A**). As the steady state protein level is determined by the rate of protein translation versus degradation, and the rate of production should be unaffected in these experiments, we reasoned that non-phosphorylatable Ame1 mutants may accumulate due to impaired protein degradation. The levels of Cse4, part of the centromeric nucleosome and a direct binding partner of AO at the inner kinetochore, have been shown to be regulated by ubiquitin-dependent proteolysis via the E3 ubiquitin ligase Psh1 (Ranjitkar et al., 2010). Levels of Gal-expressed Ame1-WT, however, remained low in a *psh1Δ* strain background, while Ame1-7A and -7E accumulated as in the wild-type background (**Supplementary Figure 2B**). This suggests that Psh1 is not involved in Ame1 level regulation under these conditions. Another E3 ubiquitin ligase complex, Ubr2/Mub1 has been shown to regulate Dsn1, which is a subunit of the Ame1 binding partner Mtw1c (Akiyoshi et al., 2013). Similar to *psh1Δ*, however, Ame1 protein levels were unaffected by the *mub1* deletion and we conclude that Ubr2/Mub1 is not involved in Ame1 level regulation under these conditions either (**Supplementary Figure 2C**).

**Figure 2:**
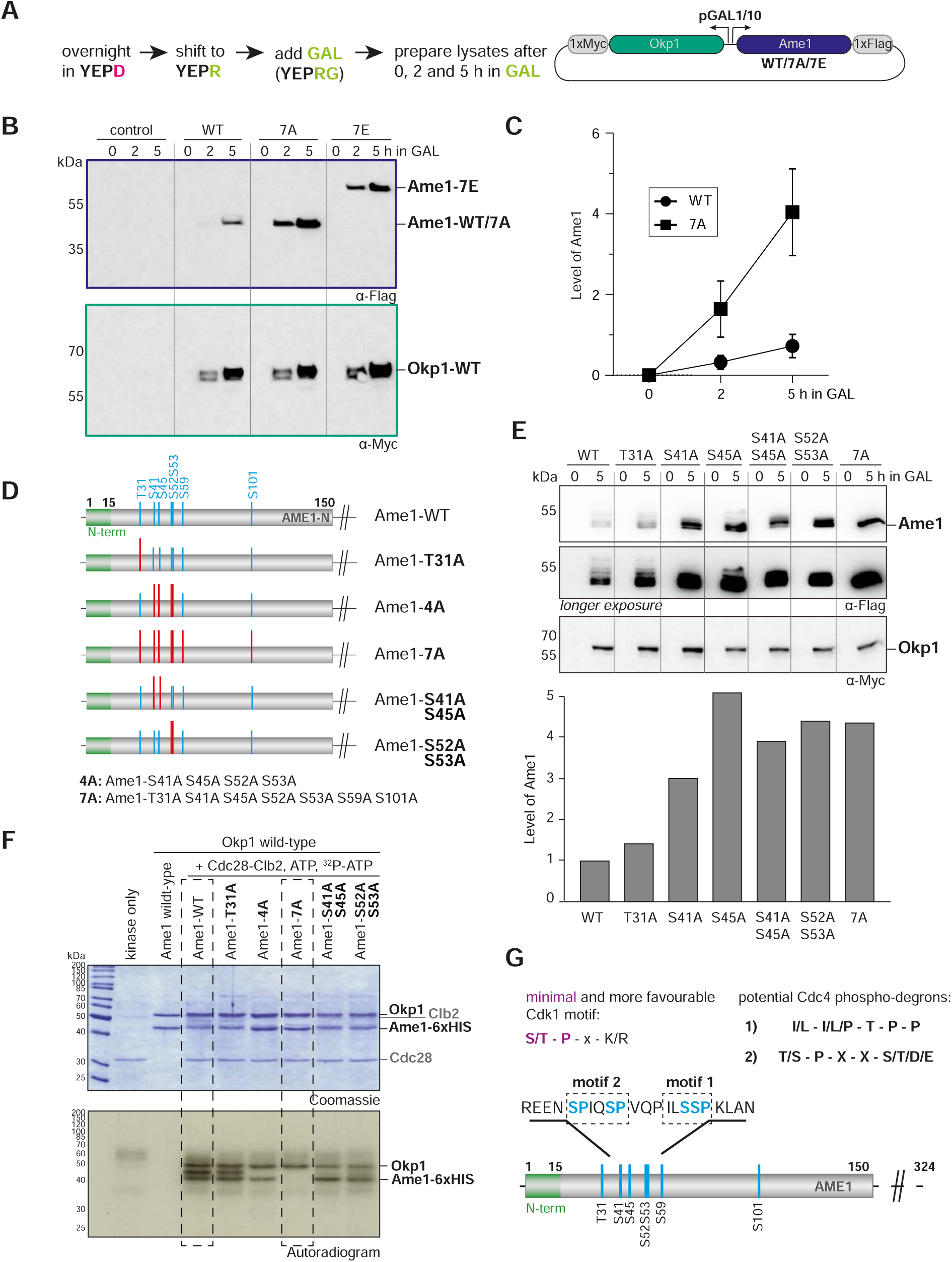
Identification of phospho-degron motifs in Ame1. **A:** Flag- and Myc-tagged versions of Ame1 and Okp1 were expressed from a two-micron plasmid under a bidirectional galactose-inducible promoter. **B:** Western Blot analysis of overexpressed Ame1-WT, 7A and 7E variants in a wild-type strain background. **C:** Quantification of protein levels of Ame1-WT and Ame1-7A after indicated times in Galactose medium. Mean values and standard error of the mean are indicated, n=7. **D:** Overview of Ame1 phospho-mutants used for overexpression studies (**E**) or in vitro kinase assays (**F**). Overexpression studies of individual Ame1 phospho-variants. Okp1 levels are stable and used for normalization for quantification of Ame1 protein levels. **F:** In vitro kinase assay of AO complexes using recombinant Cdk1-Clb2. Note reduced, or lacking phosphorylation of Ame1-4A and -7A, respectively. Also Okp1 can be phosphorylated by Cdk1-Clb2. **G:** Cdk1 target sites in Ame1 resemble two different types of phospho-degrons motifs that are recognized by the E3 ubiquitin ligase complex SCF-Cdc4.

### Identification of two phospho-degron motifs in the Ame1 N-terminus

To delineate the contribution of individual phosphorylation sites to Ame1 protein level regulation in the overexpression setting, we constructed mutants in which we prevented phosphorylation at selected sites individually or in combination (**Figure 2D**). Analysis of Ame1 protein levels after five hours of expression in the presence of Galactose showed that preventing phosphorylation on Thr31 had relatively little effect on Ame1 level when compared to the wild-type. By contrast, preventing phosphorylation at Ser41, Ser45 or Ser52/53 led to accumulation of the protein, roughly similar to preventing phosphorylation altogether in the 7A mutant (**Figure 2E**). We prepared the analogous Ame1 mutants as recombinant Ame1-Okp1 (AO) complexes for in vitro kinase assays to evaluate the contribution of these individual sites to overall Ame1 phosphorylation. Autoradiographs showed that in addition to Ame1, also Okp1 can be phosphorylated by Cdc28-Clb2 (**Figure 2F**, see also **Supplementary Figure 1 and 3**). Ame1-WT appeared as two separated phosphorylated forms after in vitro phosphorylation. The Ame1-T31A mutant displayed a similar phosphorylation pattern, while the phosphorylation of Ame1-4A was clearly decreased, with only the fast migrating Ame1 form remaining. The Ame1-7A mutant completely eliminated Cdc28-Clb2 phosphorylation in vitro. Preventing phosphorylation on either Ser41/Ser45 or Ser52/Ser53 allowed some residual phosphorylation, but clearly decreased phosphorylation compared to wild-type. We conclude that the residues responsible for Ame1 level regulation in vivo are major targets for Cdc28 phosphorylation in vitro. Further analysis confirmed that mutating the candidate Cdk1 site Ser26 in Okp1 to alanine prevented Cdc28 phosphorylation and that the Ame1-7A/Okp1-1A complex was completely refractory to Cdc28 phosphorylation (**Supplementary Figure 3**).

Posttranslational modification via phosphorylation can be mechanistically linked to the control of protein stability via the generation of so called phospho-degrons (Skowyra et al., 1997). The best studied example for this mechanism is the controlled ubiquitination and degradation of key cell cycle regulators by modular SCF complexes, using F-box proteins as readers of phosphorylated substrates (Feldman et al., 1997; Ord et al., 2019a; Ord et al., 2019b). Intriguingly, the Ame1 N-terminal sequences resembled previously described Cdc4 phospho-degrons: The Ame1 sequence surrounding Ser52 and Ser53 (motif 1) showed resemblance to a Cyclin E-type phospho-degron, comprehensively described in the context of the Cdk1 inhibitor Sic1 (Koivomagi et al., 2011; Nash et al., 2001), while the combination of Ser41/Ser45 (motif 2) resembled a di-phospho-degron with a typical +4 spacing between phosphorylated residues found for example in the Acetyltransferase Eco1 (Hao et al., 2007; Lyons et al., 2013; Lyons and Morgan, 2011) (**Figure 2G**).

### Molecular dynamics simulations predict Ame1 phospho-peptide binding to Cdc4

To evaluate candidate degron motifs in Ame1, we performed Gaussian accelerated Molecular Dynamics (GaMD) simulations of Ame1 phospho-peptide binding to the WD40 domain of Cdc4, using as template for initial coordinates the published crystal structure of a Cyclin E-derived model peptide associated with Cdc4 (Orlicky et al., 2003). The analysis of the trajectories allowed us to predict that the doubly phosphorylated Ser41/Ser45 peptide should be a good Cdc4 binder. Phospho-Ser41 establishes hydrogen bond interactions with the guanidino groups of Arg485 and Arg534 of Cdc4 while phospho-Ser45 of the peptide establishes hydrogen bond interactions with Arg467 of Cdc4 (**Figure 3A**). Further, phospho-Ser45 is engaged in additional interactions with Arg443, Ser464, and Thr465 of Cdc4 (**Figure 3B**). The peptide residues Pro42, Ile43, Glu39, and Asn40 are also involved in interactions with the protein (**Figure 3B**). Overall, the doubly phosphorylated peptide showed a strong hydrogen bond network at the binding pocket of Cdc4, highlighting the potential of this peptide as a Cdc4 binder.

**Figure 3:**
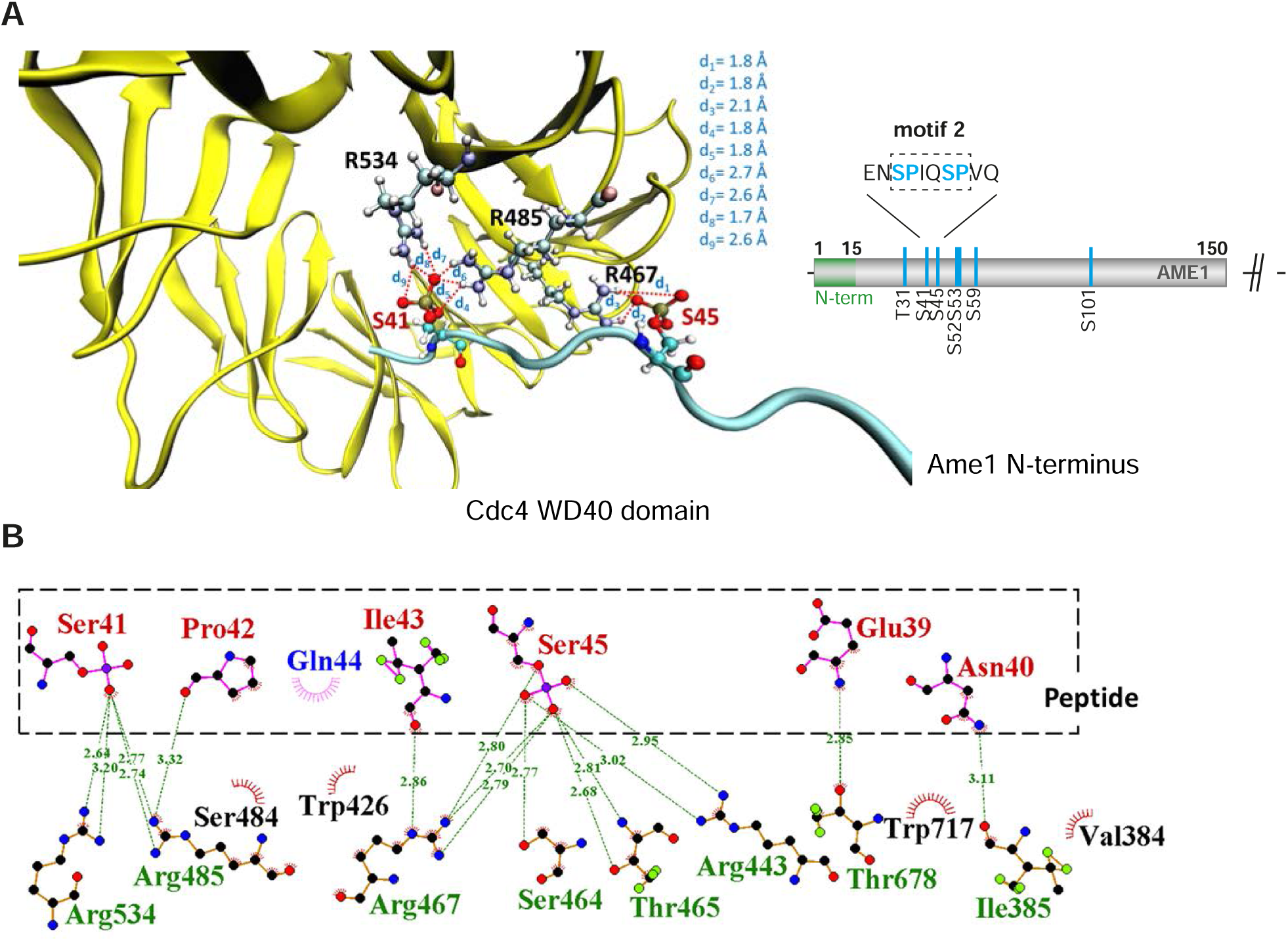
Gaussian accelerated Molecular Dynamics Simulations predict Ame1 peptide binding to Cdc4. **A:** Interactions between the conserved arginine residues of Cdc4 (yellow) and the phospho-serine residues of the doubly phosphorylated peptide (magenta). **B:** The doubly phosphorylated Ser41/Ser45 peptide and Cdc4 establish an intense hydrogen bond network involving the phosphorylated residues of the peptide and the conserved arginine residues of Cdc4, as well as other residues.

The simulations also indicated that the singly phosphorylated variants establish less interactions with Cdc4 with respect to the doubly phosphorylated Ser41/Ser45 peptide (**Supplementary Figure 4B, Supplementary Movies 1,2 and 3**). Furthermore, the protein-peptide complex involving the doubly phosphorylated peptide (**Supplementary Figure 4B, Supplementary Movies 1,2 and 3**) displayed less structural fluctuations during the trajectories compared to the simulations of Cdc4 with the monophosphorylated peptides, especially the Ser45 phosphorylated peptide. Accordingly, the doubly phosphorylated Ser41/Ser45 peptide remained attached at the binding site of the protein by establishing conserved interactions throughout the simulations, unlike the monophosphorylated peptides that displayed a dynamic behavior with less retention of their binding motifs (**Supplementary Movies 1,2 and 3**).

### The SCF Ligase with the F-box protein Cdc4 regulates Ame1-Okp1 protein levels

To test whether Ame1 level regulation is under control of SCF via phospho-degrons in vivo we used different SCF mutant alleles in the GAL overexpression setting, starting with Skp1 as a component of all modular SCF complexes. All SCF *ts* alleles were used at the permissive temperature, ensuring progression through the cell cycle. Western blotting showed that Ame1-WT expressed from the GAL promoter strongly accumulated in the *skp1-3* mutant relative to a wild-type background (**Figure 4A**). The phospho-forms of Ame1 were preserved under these conditions, showing that the *skp1-3* mutant uncouples phosphorylation of Ame1 from its degradation. Interestingly, under these conditions, also an accumulation of Okp1, expressed from the same plasmid, was apparent. Okp1 appeared in two distinctly migrating forms, possibly corresponding to phosphorylation. Combining the Ame1-7A mutation with the *skp1-3* background revealed that the Ame1-7A protein was further enriched in the *skp1-3* background compared to the wild-type strain, demonstrating that Ame1-7A can be further accumulated by compromising the SCF machinery in addition to preventing phosphorylation of Ame1 itself. We extended this analysis to mutant alleles in other SCF subunits, in particular to identify which F-box protein is responsible for Ame1 regulation. Similar to the *skp1-3* mutant, overexpressed Ame1 accumulated in mutant alleles of the Cullin subunit Cdc53, the E2 enzyme Cdc34 and the F-Box protein Cdc4. By contrast Ame1 levels remained low (or were even decreased relative to wild-type) in a deletion mutant of the cytoplasmic F-box protein Grr1 (**Figure 4B**). Interestingly, the SCF mutants also had a pronounced effect on the level of overexpressed Okp1, with particularly strong accumulation (30-fold increase) observable in the *cdc4-1* mutant. In the background of the *cdc34-2* allele, Okp1 accumulated only slightly when Ame1 was wild-type, but more strongly when phosphorylation of Ame1 was prevented (**Figure 4B**). This indicates that in the context of the Ame1-Okp1 complex, level regulation by phosphorylation may occur both in *cis* (only affecting the subunit itself) or in *trans* (affecting also an interaction partner).

**Figure 4:**
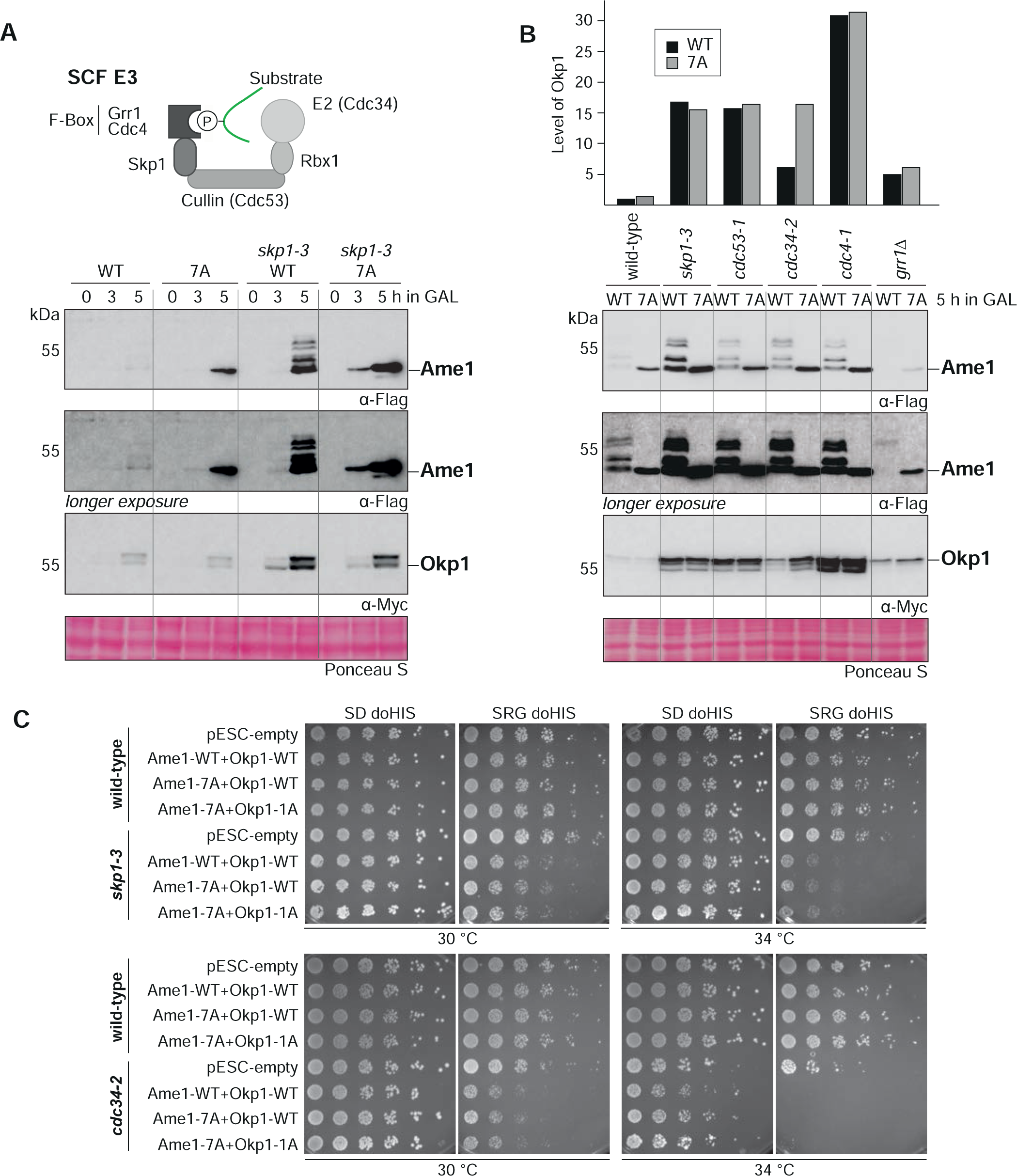
SCF-Cdc4 regulates Ame1-Okp1 protein levels in vivo. **A:** Model of substrate binding to SCF complexes. SCF is composed of Skp1, Cdc53 (Cullin), Rbx1, an F-Box protein (e.g. Cdc4 or Grr1) and here with the E2 enzyme Cdc34. Lower Panel: Expression of Ame1-WT leads to accumulation of the protein in a *skp1-3* mutant strain. **B:** Protein levels of Ame1-Okp1 in different SCF mutants. Note that Ame1 levels remain low in the *grr1Δ* mutant (cytoplasmic F-Box protein), and that Okp1 strongly accumulates in the *cdc4-1* mutant. All alleles were used at the permissive temperature of 30 °C. **C:** Serial dilution assay of overexpressed Ame1-Okp1 variants in wild-type or SCF mutant strain backgrounds (*skp1-3* or *cdc34-2*). Plates were photographed after two days at the indicated temperature.

We tested the effect of Ame1-Okp1 expression from a GAL promoter in a serial dilution assay. In a wild-type strain background, AO overexpression was tolerated well. In a *skp1-3* mutant background however, overexpression of AO, either in wild-type form or with Cdk1 sites mutated to alanine, compromised growth at 30 °C and 34 °C (**Figure 4C**). Similar results were obtained for the *cdc34-2* mutant background, in which overexpression of AO already greatly impaired growth at 30 °C. These effects are consistent with AO being physiological substrates of the SCF machinery and they show that accumulation of AO can negatively impact cell growth.

### Increasing degron strength in the Ame1 N-terminus partially suppresses a *cdc4* mutant

To further study the regulation of Ame1-Okp1 by phospho-degrons we converted the motif 1 degron (ILSSP) of Ame1 into a stronger Cdc4 phospho-degron, which was shown in the context of the Cdk1 inhibitor Sic1 to provide the highest binding affinity for Cdc4 (Ame1-CPD^ILTPP^ (Nash et al., 2001)). GaMD simulations were performed to study the binding of this Ame1-derived peptide featuring a phosphorylated threonine at position 52 to Cdc4 (sequence VQPIL**TP**PKL, as in the Ser-phosphorylated peptides, 3 replicas, 100 ns each). The analysis of the resulting trajectories indicated that the peptide establishes conserved interactions at the binding pocket of the protein and remains bound (**Supplementary Movie 4**, **Supplementary Figure 4E**). A representative snapshot of the most populated cluster of structures, along with the overall population of that cluster is shown in **Supplementary Figure 4C**. The phosphorylated threonine Thr52 establishes a strong hydrogen bond network with the conserved arginine residues of Cdc4: Arg467, Arg485, and Arg534 (**Figure 5A**). Further, Leu56 of the peptide interacts with Arg443 and Thr465 of Cdc4, while the peptide residues Val47, Pro53, and Pro54 also interact with protein residues (**Supplementary Figure 4D**). Additionally, several van der Waals contacts are established between the peptide (through Pro49, Leu51, Lys55, and Ile50) and Cdc4 (involving Leu637, Thr677, Ile676, Trp717, Ser464, Tyr574, and Gly636), indicating an optimal fit of the peptide at the protein binding site. Overall, this strong network of peptide-protein interactions indicates that the VQPIL**TP**PKL peptide is predicted to be a potent Cdc4 binder, even more than the doubly phosphorylated Ser41/Ser45 peptide.

**Figure 5:**
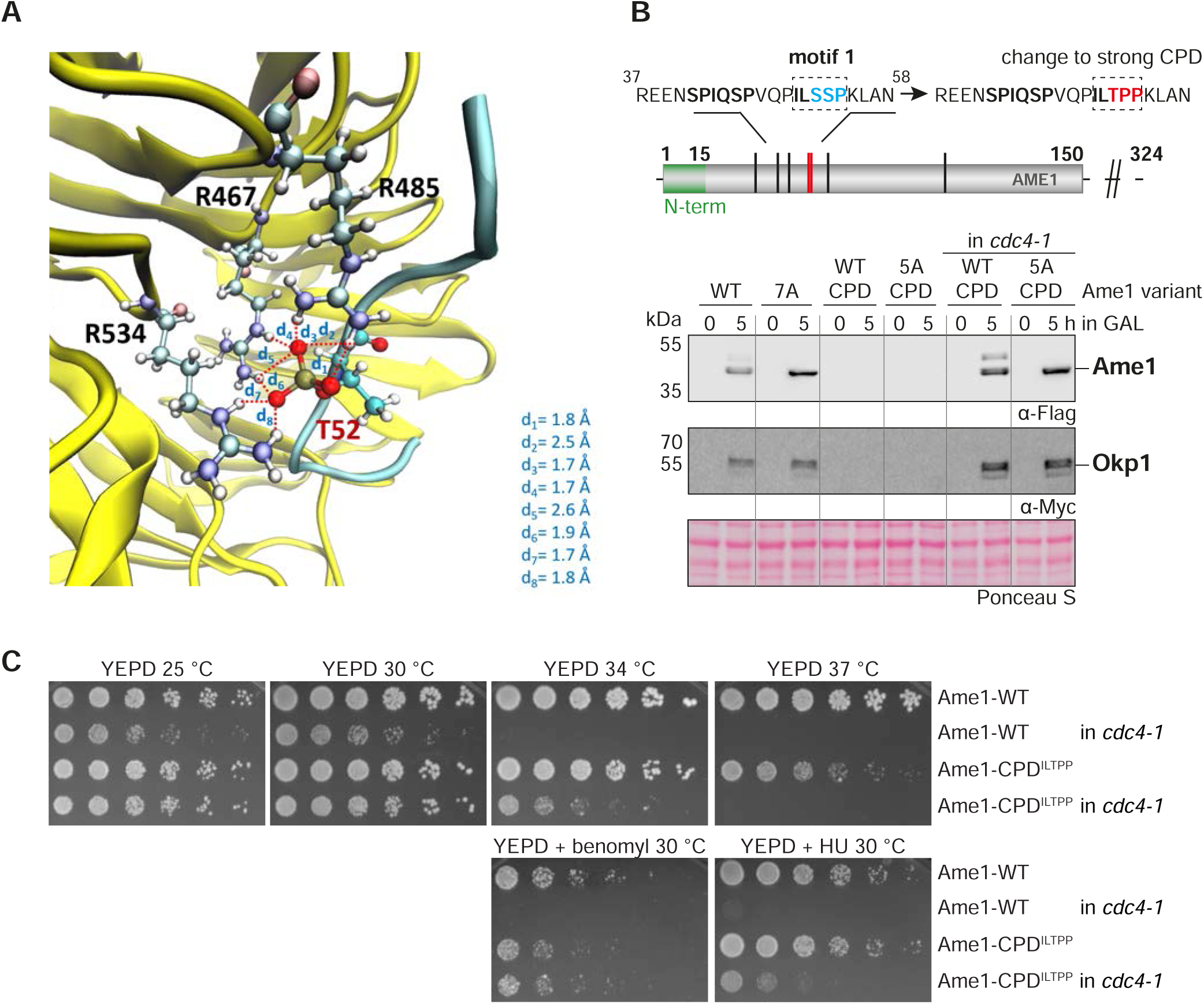
Tuning degron strength in the Ame1 N-terminus suppresses a *cdc4* mutant. **A:** The threonine phosphorylated peptide (VQPIL**TP**PKL, magenta) establishes a strong network of conserved interactions involving its phosphorylated threonine and the conserved arginine residues of Cdc4 (yellow). This binding is further stabilized by several protein-peptide interactions (**Supplementary Figure 4D**). **B:** Changing the phospho-degron motif 1 into a strong Cdc4-degron sequence (ILSSP to ILTPP) leads to a loss of detectable Ame1-CPD in the overexpression system. Also, note that Okp1 is not detectable anymore. The *cdc4-1* mutant background stabilizes Ame1-CPD^ILTPP^ and Okp1. The *cdc4-1* allele was used at the permissive temperature of 30 °C**. C:** Serial dilution assay of Ame1-WT or Ame1-CPD in a wild-type or *cdc4-1* mutant strain background. Plates were photographed after three days of incubation at the indicated temperature. Note that Ame1-CPD partially suppresses the growth defect of *cdc4-1* at 34 °C, or in the presence of benomyl or hydroxyurea.

Next, we evaluated the Ame1-CPD^ILTPP^ mutation in the GAL overexpression setting. Strikingly, neither Ame1 nor Okp1 protein was detectable by western blotting upon overexpression under these conditions and preventing phosphorylation on the five remaining sites (Ame1-5A-CPD^ILTPP^) did not stabilize the protein (**Figure 5B**). If the strong CPD indeed exerts its effect via Cdc4-dependent recognition, then the protein levels of Ame1-CPD^ILTPP^ should be restored to wild-type in an SCF mutant. Combining the Ame1-CPD^ILTPP^ allele with the *cdc4-1* mutant demonstrated that this is indeed the case: The Ame1-CPD^ILTPP^ mutant and also Okp1 were detectable and displayed similar levels as the wild-type proteins in a *cdc4-1* background (**Figure 5B**). These experiments provide evidence that protein levels of Ame1 and Okp1 are regulated by activation of phospho-degrons in an SCF-Cdc4 dependent manner. We also tested the effect of changing motif 1 into a strong CPD in the context of endogenous Ame1. Interestingly, Ame1-CPD^ILTPP^ expressed as the sole copy of Ame1 yielded a viable strain which showed slight temperature-sensitivity at 37 °C and increased sensitivity to benomyl (**Figure 5C**). Upon combination with a *cdc4-1* mutant, however, Ame1-CPD^ILTPP^ was able to partially suppress the growth defect of *cdc4-1* at 34 °C and improved growth in the presence of hydroxyurea and benomyl. Thus, whereas the Ame1-CPD^ILTPP^ allele reduced fitness in a wild-type strain background, it conferred a growth advantage to *cdc4-1* mutants.

### Phospho-degrons of endogenous Ame1 are activated in a cell-cycle dependent manner

In the experiments described above, cells were challenged with increased levels of Ame1 following expression from a GAL promoter. How does this relate to the regulation of endogenous Ame1? To test this, we constructed Ame1 mutants expressed from their endogenous promoter. To simplify the complex phosphorylation pattern of wild-type Ame1, we generated alanine mutants that either allowed phosphorylation of the motifs 1 and 2, but prevented phosphorylation of the remaining sites (Ame1-3A, CPD only), or, conversely, prevented motif 1 and 2 phosphorylation, but allowed the remaining sites to be phosphorylated (Ame1-4A, CPD null). Western blotting showed that motif 1/2 phosphorylation of endogenous Ame1 was cell cycle dependent (**Figure 6A**). S-Phase arrested cells displayed a single slowly migrating Ame1 form in addition to unmodified Ame1, while M-Phase arrested cells were maximally phosphorylated with two slowly migrating forms becoming apparent. In the Ame1-4A mutant (CPD null), all slowly migrating forms were eliminated (**Figure 6A**). We followed Ame1 phosphorylation over the course of the cell cycle after release from alpha-factor. Consistent with the analysis of the arrests, we observed that motif 1/2 phosphorylation occurred in a step-wise manner with fully phosphorylated forms appearing 30 minutes into the cell cycle. FACS analysis demonstrated that by this time, cells had completed replication and were in M-Phase with a 2C DNA content (**Figure 6B**). Strikingly, phosphorylated Ame1 forms gradually diminished between minute 30 and 60 and only unmodified Ame1 and a single slowly migrating form remained (**Figure 6B**). This indicates that degron motifs on authentic Ame1 are phosphorylated in a cell cycle dependent manner and lead to disappearance of the phosphorylated forms in M-Phase. We additionally analyzed Pds1 degradation kinetics in different Ame1 mutants over the cell cycle. Ame1-3A phosphorylation was highly reproducible, while Ame1-4A eliminated all phosphorylation events observable by shift (**Figure 6C, 6D**). Interestingly, Ame1 mutants in which only phosphorylation on motif 1 was allowed (Ame1-5A1) or only on motif 2 (Ame1-5A2) also largely eliminated the characteristic stepwise phosphorylation of Ame1-3A (**Supplementary Figure 5A** and **B**), showing that full phosphorylation is mutually dependent on the presence of both motif 1 and motif 2. Compared to wild-type, neither of the mutants led to a strong delay in the cell cycle as judged by Pds1 degradation kinetics or FACS analysis.

**Figure 6:**
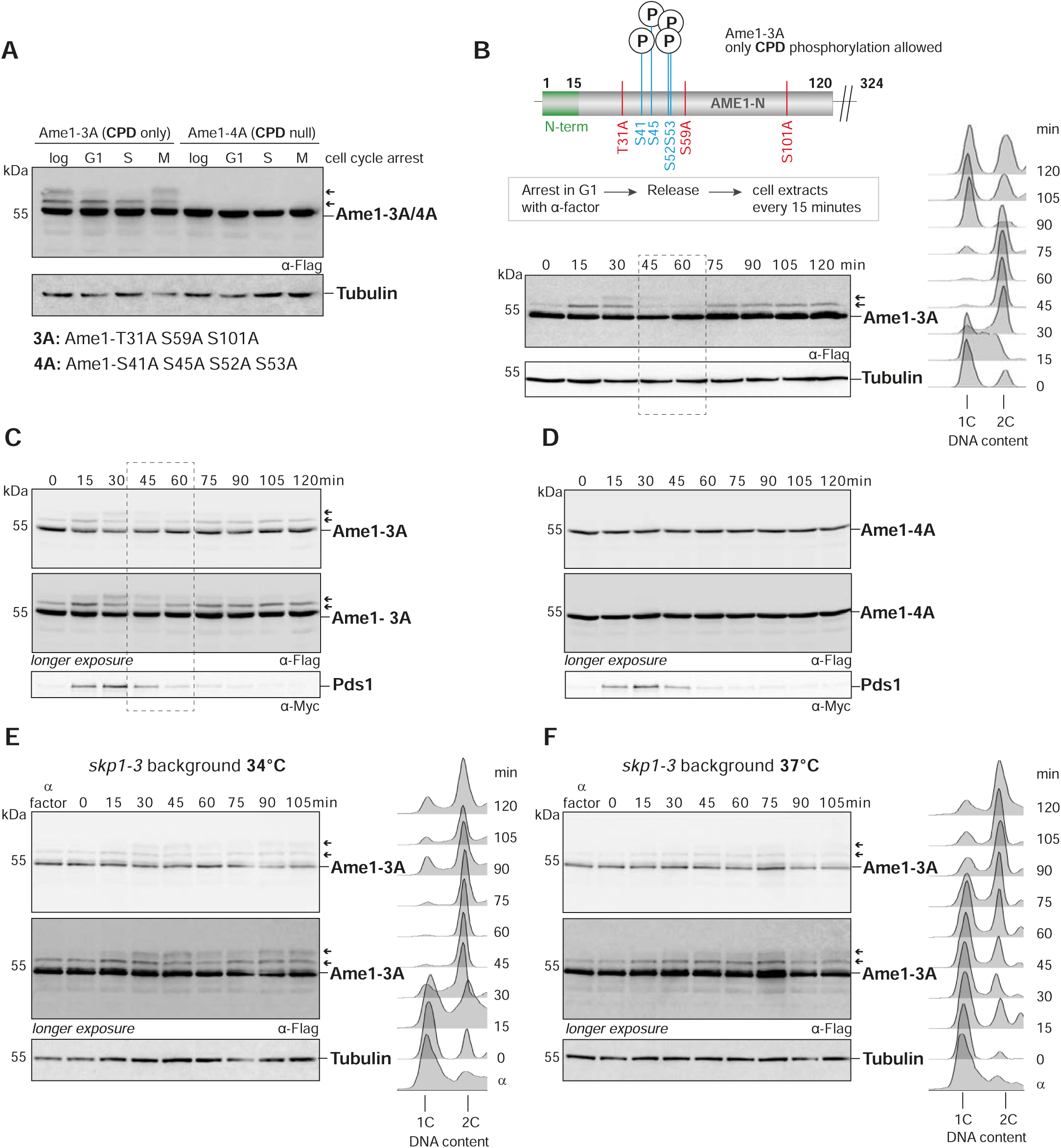
Analysis of endogenous Ame1 phospho-mutants over the cell cycle. **A:** Ame1-variants (Ame1-3A, CPD only, allowing phosphorylation at degron motifs 1 and 2 or eliminating it, Ame1-4A, CPD null) were expressed from the endogenous promoter as the sole copy and phosphorylation was analyzed in different cell cycle arrests. Ame1-3A shows one slowly migrating form in S-Phase and two in M-Phase, whereas Ame1-4A eliminates all slowly migrating forms. **B**: Ame1-3A was released from an alpha-factor arrest and phosphorylation was analyzed by western blotting. Right panel: DNA content analysis by FACS. Phosphorylation is maximal after 30 minutes when cells have completed S-Phase (right: FACS analysis 15+30 minutes) and phosphorylated forms disappear when cells are in mitosis (45+60 minutes). **C-D:** Cell cycle analysis of Ame1-3A (**C**) and Ame1-4A (**D**). **E:** Analysis of Ame1-3A in the *skp1-3* mutant at 34 °C (semi-permissive). Note that phosphorylation at motif 1+2 persists in the mutant and cells remain in mitosis with 2C DNA content. **F:** Analysis of Ame1-3A in the *skp1-3* mutant at 37 °C (restrictive). Note that under these cells are delayed to complete replication and mainly a single phospho-form of Ame1 is found.

We next combined the Ame1-3A allele with a *skp1-3* mutant and analyzed progression through the cell cycle. We found that at the semi-permissive temperature of 34 °C *skp1-3* cells completed replication with nearly wild-type kinetics and accumulated phosphorylated Ame1. Strikingly, in contrast to wild-type cells, phosphorylated Ame1 persisted and the cells remained in mitosis with a 2C DNA content (**Figure 6E**). When assayed at the restrictive temperature of 37 °C *skp1-3* cells only slowly progressed to 2C DNA content, indicating they had problems initiating and completing replication. In this situation, Ame1-3A showed only a singly phosphorylated form in western blotting and was only slowly phosphorylated to full extent (**Figure 6F**). These experiments show that use of different temperatures allows a separation-of-function of the *skp1-3* allele: At 34 °C specifically the mitotic function of Skp1 is compromised and this is accompanied by the inability to eliminate Ame1 phosphorylated at the motifs 1 and 2.

### Binding of the Mtw1 complex shields Ame1 phospho-degrons from Cdk1 phosphorylation

Our analysis described above suggests that Cdk1 phosphorylation targets Ame1 for degradation during mitosis via SCF. This seems counterintuitive, given the critical function of the kinetochore in mitosis. The Western Blot analysis also indicates, however, that only a subset of endogenous Ame1 is subjected to phosphorylation. Since the phosphorylation cluster is close to the Mtw1 binding site, we asked how binding of the Mtw1c to Ame1 affects phosphorylation by Cdc28-Clb2 in vitro (**Figure 7A**). Analysis of in vitro kinase assays revealed that while Ame1 was heavily phosphorylated in isolation, inclusion of the Mtw1 complex strongly reduced the incorporation of phosphate into Ame1 under identical conditions (**Figure 7B**). This reduction was not due to competing phosphorylation of the Mtw1c, as only the Dsn1 subunit seemed to be a minor substrate of Cdc28-Clb2. We conclude that free AO is the preferred substrate for Cdk1 phosphorylation. This could explain how kinetochore-bound Ame1 is protected from phosphorylation and subsequent degradation in mitosis.

**Figure 7:**
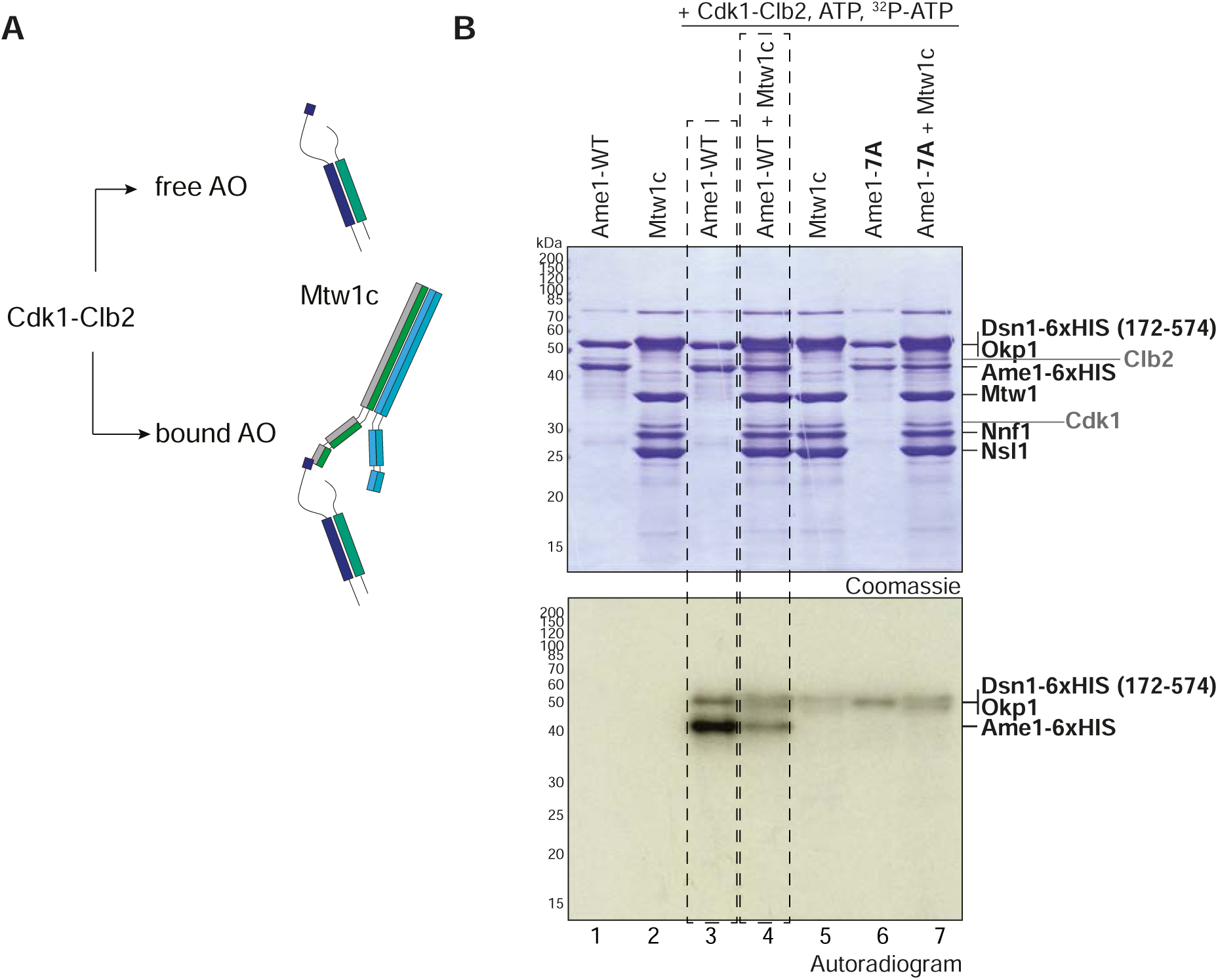
Mtw1c binding shields the Ame1 phospho-degron from Cdk1 phosphorylation. **A:** Scheme of the kinase assay. Recombinant AO with Ame1-WT or Ame1-7A is used either alone or in combination with its binding partner Mtw1c. **B**: In vitro kinase assay of Ame1-Okp1c alone or preincubated with Mtw1c and Cdk1-Clb2 shows decreased phosphorylation of Ame1-WT-Okp1c when bound to Mtw1c (lanes 3+4). Phosphorylation of Okp1 is overlapping with phosphorylation of Dsn1 (lanes 4+5+7).

### Overexpression of COMA is toxic to cells

The observation that non-phosphorylatable Ame1 was further stabilized in SCF mutants (see **Figure 3 A, B**) and that in contrast to the *skp1-3* mutant at 34 °C, the phospho-null allele failed to delay cells in mitosis, suggested that additional SCF substrates at the kinetochore may exist. Ame1-Okp1 associates with Ctf19-Mcm21 (CM) to form the COMA complex, and our phosphorylation analysis identified Mcm21 as a Cdk1 substrate with a well conserved SP site at Ser139 (**Figure 8A**). Interestingly, both in vivo and in vitro phosphorylation analysis also found the preceding Thr138 to be phosphorylated.

**Figure 8:**
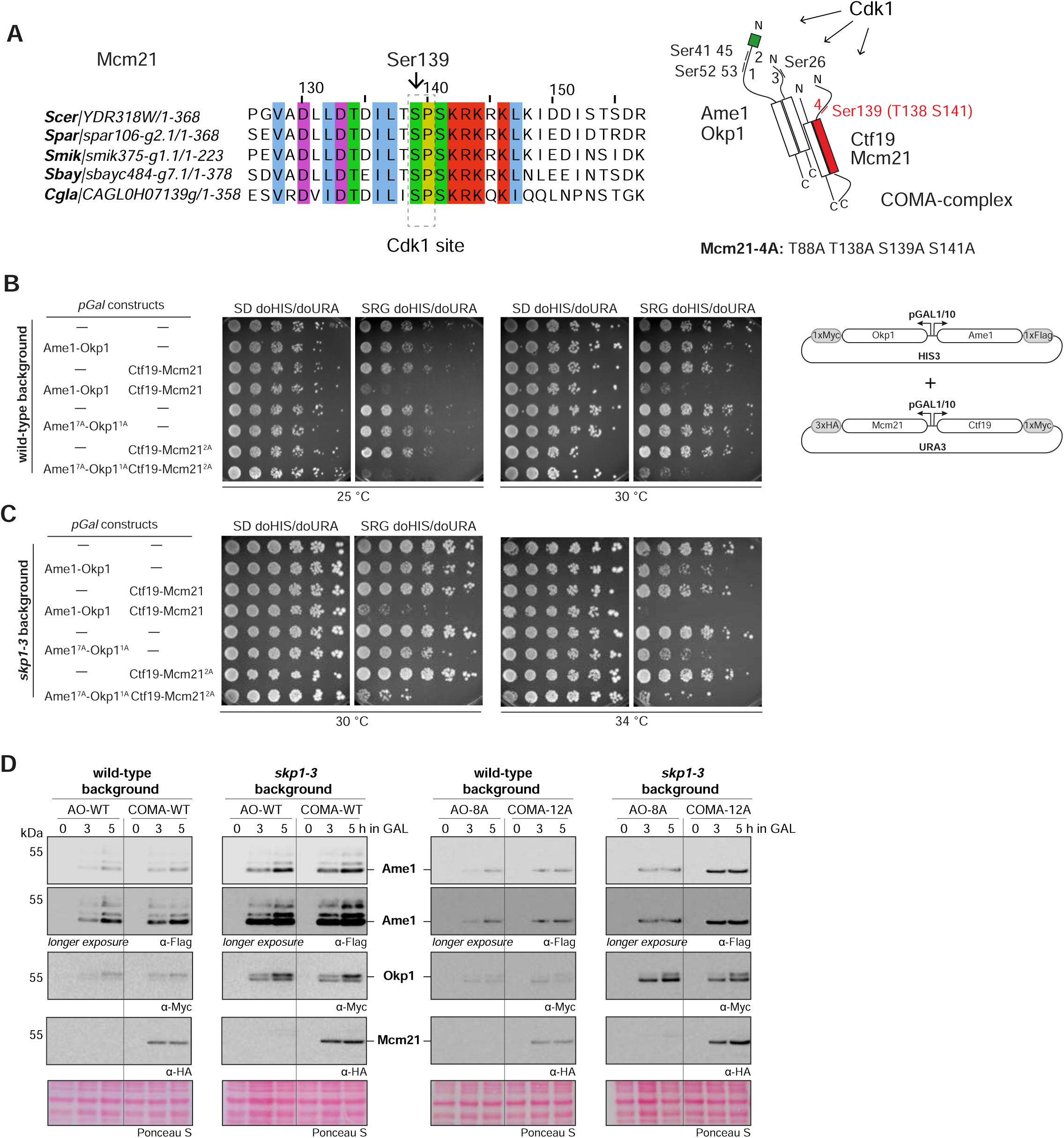
Elevated levels of COMA subunits impair cell growth. **A.** Multiple sequence alignment of Mcm21 proteins from different species, showing conservation of the Ser139 phosphorylation site and surrounding residues. Right panel: schematic overview of Cdk1 phosphorylation sites and degron motifs in COMA complex subunits. **B:** Serial dilution assay of overexpressed Ame1-Okp1, Ctf19-Mcm21 or COMA in a wild-type strain background. SD: Glucose-containing minimal medium, SRG: Galactose-containing minimal medium. **C:** Serial dilution assay of overexpressed Ame1-Okp1, Ctf19-Mcm21 or COMA in a *skp1-3* SCF mutant background. **D:** Western Blot analysis of overexpressed AO or COMA in wild-type or phospho-mutant (Cdk1 sites) at indicated times after shift to Galactose-containing medium.

We used the GAL-induced overexpression setting to address the role of AO versus COMA phosphorylation in more detail. We first tested the effect of overexpressing AO or CM individually or together as the full COMA complex in a wild-type strain (**Figure 8B**). While expression of AO or CM individually was tolerated well, overexpression of COMA severely compromised growth (**Figure 8B**). Similar results were obtained in the sensitized *skp1-3* background (**Figure 8C**). As demonstrated before (**Figure 4C**), overexpression of AO compromised growth of the *skp1-3* mutant at 34 °C, however expression of COMA had a much stronger effect. Overexpression of non-phoshorylatable AO in *skp1-3* yielded a more pronounced growth phenotype compared to wild-type AO, this was most noticeable at 30 °C (**Figure 8C**). Western Blot analysis indicated that non-Cdk1-phosphorylatable COMA accumulated more strongly than the corresponding non-phosphorylatable Ame1, especially in the *skp1-3* mutant background (**Figure 8D**). In addition this analysis demonstrates that mutation of Ser26 on Okp1 does not fully eliminate the slowly migrating form, suggesting that in addition to Cdk1, other kinases must contribute to the regulation of Okp1.

## Discussion

This study reveals novel aspects of phospho-regulation at the budding yeast inner kinetochore. We show that an important function of Cdk1 phosphorylation is to generate phospho-degron motifs on selected inner kinetochore subunits, including the essential COMA subunit Ame1, which are then recognized by the conserved ubiquitin ligase complex SCF with its phospho-adapter Cdc4. We note that Ame1 and Mcm21 peptides were also identified in a large-scale proteomic study geared towards enriching peptides simultaneously regulated by phosphorylation and ubiquitination (Swaney et al., 2013). In this context, ubiquitination of COMA subunits was detected for Okp1 (on residue Lys57) and Mcm21 (on residue Lys229).

While Cdk1 has been thought to promote kinetochore assembly in most contexts investigated so far, our study indicates that it can also act as a negative regulator of kinetochore assembly by targeting subunits for ubiquitination and subsequent degradation by the proteasome. This seems counterintuitive at first, given that kinetochores perform their essential role in segregating sister chromatids during mitosis. There are, however, important aspects in which the inner kinetochore subunit Ame1 differs from previously studied SCF substrates: While for example the Cdk1 inhibitor Sic1 is fully degraded at the G1-S transition to allow replication initiation, our experiments indicate that only a subset of Ame1 is phosphorylated and subjected to the SCF-dependent pathway. Our biochemical experiments furthermore suggest that the pool of Ame1 regulated by this mechanism corresponds to molecules that are not bound to their binding partner within the kinetochore, the Mtw1c. Such excess Ame1 subunits were also present in our GAL-induced overexpression setting, in which we initially characterized the SCF-dependent regulation of Ame1. While this experiment creates an artificial situation in which the cell is challenged with an increased level of Ame1, this scenario likely also applies to the natural kinetochore assembly process. To ensure effective assembly during S-Phase, free kinetochore subcomplexes must be present in excess amounts, otherwise they would become limiting for assembly and prevent the effective formation of a new kinetochore. On the other hand, excess free subcomplexes could favor ectopic assembly which would lead to genetic instability. As shown in **Figure 7**, Mtw1c binding of Ame1-Okp1 subcomplexes shields degron phosphorylation. This Mtw1c binding sensitive phosphorylation could ensure that only free, unused subcomplexes are removed by degradation (**Figure 9A**). From a structural standpoint, kinetochore subcomplexes typically combine relatively short, structured segments (often coiled-coil domains) with large unstructured domains that are the preferred targets of phosphorylation. In this context, phospho-degrons could be ideally suited as assembly-sensors for kinetochores since they allow to distinguish excess subunits from properly assembled ones. By placing individually weak degron signals on separate subunits, the cell may allow COMA assembly from AO and CM complexes, while phosphorylation of the assembled COMA then creates stronger composite binding sites for Cdc4 (**Figure 9A**).

**Figure 9:**
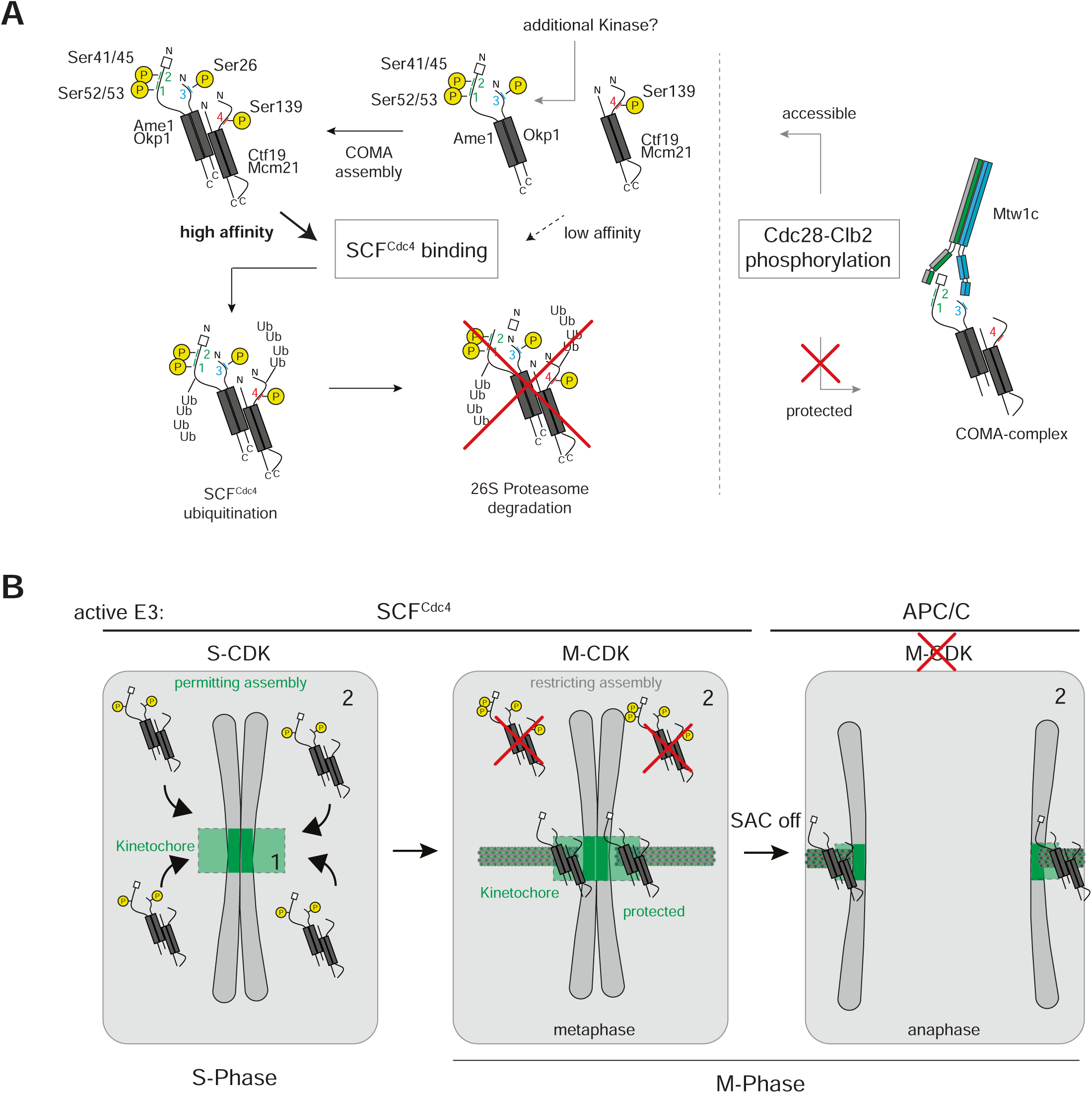
Model for SCF-mediated regulation of COMA assembly at the budding yeast kinetochore. **A.** Scheme illustrating stepwise phosphorylation of degron motifs on Ame1 and Mcm21 by Cdk1 and protection from phosphorylation within the kinetochore. In addition to Cdk1, other kinases might be involved in addition, in particular in the phoshphorylation of Okp1 **B.** Cell cycle regulation of COMA complex stability, for details see discussion. In S-Phase, COMA is only partially phosphorylated, allowing assembly at the kinetochore. In M-Phase, free COMA is fully phosphorylated and targeted for degradation, while kinetochore-bound COMA is protected.

The observation that even placing the strong CPD^ILTPP^ degron into Ame1 does not eliminate the endogenous protein fully, is consistent with the idea that at least some budding yeast kinetochores are always assembled and that therefore a significant fraction of Ame1 is continuously protected from phosphorylation and subsequent degradation. Elimination of those COMA molecules that are not bound to Mtw1c might be important for multiple reasons: Increased levels of COMA may be dangerous because they could facilitate the formation of ectopic kinetochores. Indeed, we show that GAL-based overexpression of all four subunits of the COMA complex is toxic for cells. COMA might be an especially important target for regulation in this regard, because it is near the top of the assembly hierarchy and contacts multiple other inner kinetochore subunits (Yan et al., 2019). It also contains DNA-binding elements, such as AT hooks, that may allow it to bind to chromosome arms when not properly targeted to centromeres.

In summary, we propose the following model for COMA phospho-regulation by Cdc28^Cdk1^ (**Figure 9B**): Initial phosphorylation on Ame1 motifs 1 and 2 starts in S-Phase but is not complete before M-Phase. This ensures that sufficient free subcomplexes are available for kinetochore assembly. In parallel, the observation that individual degrons on AO or CM are weak, permits COMA assembly from its subcomplexes. In M-Phase, full degron phosphorylation destabilizes assembled COMA complexes, unless they are bound to the Mtw1 complex, which shields the degrons. The proper timing of phosphorylation is critical in this model. Premature phosphorylation of COMA would target it for destruction too early, likely compromising kinetochore assembly in S-Phase. This could be the reason, why the Ame1 phospho-sites are only gradually phosphorylated in vivo. Conversely, placing the stronger CPD^ILTPP^ phospho-degron into Ame1 compromised growth in otherwise wild-type cells, but partially rescued the growth phenotypes of *cdc4-1* mutants. This argues that indeed key mitotic SCF substrates reside at the inner kinetochore. Notably, the Histone H3 variant Cse4 has recently been reported to be an SCF substrate (Au et al., 2020).

While our experiments strongly implicate Ame1 as a Cdk1 and SCF substrate, they also show that the Ame1-7A mutant effectively prevents phosphorylation in vivo and in vitro, but does not induce a strong mitotic delay such as the *skp1-3* mutant. We speculate that the Ame1-7A mutant is not sufficient to fully prevent SCF regulation of the inner kinetochore, and additional phospho-targets must exist. The molecular requirements for the recognition of phosphorylated substrates may be complex in the case of multi-protein complexes such as COMA. Here, degron sequences located on multiple subunits and phosphorylated in different combinations might be required for effective binding by Cdc4. This could involve both non-consensus sites phosphorylated by Cdk1, and also kinases in addition to Cdk1, as is the case for multiple other SCF substrates (Faustova et al., 2021; Koivomagi et al., 2011; Ord et al., 2019a). An additional key substrate may be Okp1, and some Cdk1-dependent phosphorylation sites (e.g. Ser70) have been reported (Holt et al., 2009), which are not direct Cdk1 sites. Candidate kinases that may be involved include Cdc5/Plk1 and indeed Ame1 Ser52/Ser53 constitute candidate Polo-box binding sites and human CENP-U has been established as a kinetochore receptor for Plk1 (Singh et al., 2021). Biochemical reconstitution of the ubiquitination reaction with purified components in vitro can shed light on the molecular requirements and mechanisms of CCAN phosphorylation and ubiquitination by SCF in the future.

Previous work has defined the role of SCF-Cdc4 at the G1-S transition, but has also shown that Cdc4, as well as Cdc34 or Cdc53, remain essential genes in *sic1Δ* mutants (Schwob et al., 1994), demonstrating that there must be additional key substrates. Furthermore, SCF mutants that complete replication, arrest at G2-M with a short spindle and an activated mitotic checkpoint (Goh and Surana, 1999; Schwob et al., 1994) (our unpublished observations). The relevant substrates for this mitotic function of SCF-Cdc4, however, have so far remained elusive. Our experiments indicate that Ctf19^CCAN^ subunits are cell-cycle dependent substrates of SCF-Cdc4 and open up the possibility that regulation of kinetochores by SCF makes a critical contribution to the metaphase-anaphase transition. They also reveal that in addition to the well-established relationship between kinetochores and the E3 ligase APC/C, which regulates the metaphase-anaphase transition via the Mitotic Checkpoint Complex, kinetochores are also molecularly linked to SCF, the other major RING-type E3 ligase complex that regulates the eukaryotic cell cycle.

## Materials and Methods

### Expression and purification of recombinant proteins (AO, Mtw1c, MN, Cdc28-Clb2)

Expression constructs for kinetochore proteins AO, Mtw1c and Mtw1-Nnf1 (MN) and the kinase complex Cdc28-Clb2 used in this study were created by amplification of the DNA for the respective genes from yeast genomic DNA and cloning into pETDuet-1, pET3aTr/pST39, pST44 plasmids (bacterial expression) following the protocol for restriction free cloning or pESC two-micron plasmids (yeast expression) using classical cloning methods. Restriction free cloning was also used to produce vectors encoding phospho-eliminating mutants in Ame1. Site directed mutagenesis (Agilent technologies) was applied for introduction of amino acid substitutions. A list of all vectors used for protein production and purification in bacteria or insect cells can be found in **Table 3.**

Following conditions were used for protein production and purification unless indicated otherwise: Competent bacterial cells were grown at 37 °C until OD_600_ of 0.6 and subsequently induced with 0.5 mM IPTG. Expressions were conducted overnight at 18 °C-20 °C. Expression of Mtw1c and MN constructs was performed in BL21 (DE3; Novagen) and AO was expressed in Rosetta 2(pLys) (DE3) cells (EMD Millipore). Cdc28-Clb2 was overexpressed in the protease-deficient yeast strain DDY1810 overnight at 30 °C using 2 % galactose. Lysis, wash, and elution buffers as well as chromatography steps varied for the different protein complexes and are described in the individual sections. The Poly-Histidine fusion proteins were isolated with HisTrap HP 5 ml columns. The kinase complex was isolated using IgG sepharose 6 Fast Flow and Calmodulin sepharose 4B affinity resin (both GE Healthcare). In the last step of all bacterially expressed proteins, the protein was passed over a size-exclusion chromatography (SEC) column that was appropriate for the proteins size while elution of proteins was measured by absorbance at 280 nm using an Äkta FPLC System connected to a Windows based laptop running the UNICORN control software. All proteins were concentrated to the desired concentration except the kinase complex and flash frozen in liquid nitrogen before stored at −80 °C until usage.

#### Mtw1c

The Mtw1 complex (Mtw1c) used in this study includes an N-terminal 171 amino acid truncation of the Dsn1 subunit and an N-terminal 6xHis tag at the same protein. Lysis buffer for Mtw1c was 50 mM Tris-HCl, pH 7.5, 500 mM NaCl, 30 mM imidazole, 10 % glycerol, and 5 mM β-Mercaptoethanol as described previously (Maskell et al., 2010). Lysates were loaded onto a HisTrap HP 5 ml column pre-equlibrated in lysis buffer, washed with 30 column volumes (CV) lysis buffer and eluted with 30 mM Tris, pH 8.5, 80 mM NaCl, 10 % glycerol, 5 mM β-Mercaptoethanol and 250 mM imidazole. Subsequently, proteins were directly loaded onto anion exchange chromatography (HiTrap Q HP 5ml column; GE Healthcare). The column was equilibrated with 4 CV gradient wash from 0–20 % buffer B, followed by 15 CV wash of 20 % buffer B. The chromatography was performed with a gradient consisting of buffer (30 mM Tris HCl, pH 8.5, and 5 % glycerol from 80 mM [A] to 1 M NaCl [B]) using a flow rate of 1 ml/min and an elution volume of 40 CV. The protein is eluted at a buffer B ratio of about 40 %. Fractions containing the Mtw1c were concentrated using a Vivaspin 20 ultracentrifugal unit MWCO 10000 Dalton (Satorius) and loaded to a HiLoad Superdex 200 16/600 pg (GE Healthcare) equilibrated in 30 mM HEPES, pH 7.5, 250 mM NaCl, 5 % Glycerol and 2 mM TCEP.

#### Mtw1/Nnf1

In the Mtw1/Nnf1 dimer used in this study a 6xHis tag was introduced N-terminal of Nnf1. Purification was performed according to the protocol for Mtw1c described here, except that the dimer did not bind to the HiTrap Q HP 5 ml column. Therefore, the flowthrough of the column was used for concentration and subsequent loading to the Superdex 200 10/300 GL column.

#### AO

Bacterial pellets expressing the AO complex (AO) were resuspended in lysis buffer (30 mM HEPES, pH 7.5, 30 mM imidazole (A) or 1 M imidazole (B), 600 mM NaCl, and 5 mM β-Mercaptoethanol). AO loaded on the HisTrap HP 5 ml column was washed with 30 CV lysis buffer before eluted with 300 mM imidazole in the otherwise same buffer or using an imidazole gradient from 0 % to 60 %. AO eluted between 150 mM and 300 mM imidazole. Subsequently, the complex was purified over gel filtration Superdex 200 10/300 GL or HiLoad Superdex 200 16/600 pg columns (GE-Healthcare) in 30 mM HEPES, pH 7.5, 250 mM NaCl, 5 % Glycerol and 2 mM TCEP. The complex eluted in two peaks from the column, of which the later eluting peak was used for subsequent experiments. All used amino acid substitutions were purified in the same way as described above.

#### Cdc28-Clb2

Yeast cells were incubated in minimal medium under constant selective pressure (SD or S-RG doURA). Expression was induced by adding 2 % galactose to S-R doURA medium containing 2 % raffinose and additionally 10x YEP solution to boost cell growth. Yeast pellets were flash frozen in liquid nitrogen as droplets and grind into powder using a freezer mill (SPEX SamplePrep 6870). Powder was lysed in 1x HYMAN buffer (50 mM bis-Tris propane-HCl pH 7.0, 100 mM KCl, 5 mM EDTA, 5 mM EGTA, 10% glycerol; additional added protease inhibitors (Pierce Thermo Fisher, Calbiochem and 1 mM PMSF)) with 1% Triton X-100 using sonification. The lysate after high-speed centrifugation was incubated with pre-equilibrated IgG Sepharose 6 Fast Flow affinity resin (washed in TST (50 mM Tris-HCl pH 7.4, 150 mM NaCl, 0.1 % Tween 20) and NH_4_OAc pH 3.4) for 4 hours at 4 °C and the salt concentration was increased to 300 mM KCl. Beads were washed with 50 bead volumes 1x HYMAN buffer and proteins were eluted with TEV-protease overnight at 4 °C. TEV-Eluate (with increase CaCl_2_) was further incubated using preequilibrated Calmodulin Sepharose 4B affinity resin in Calmodulin binding buffer (25 mM Tris-HCl pH 8.0, 150 mM NaCl, 0.02 % NP-40, 1 mM MgCl_2_, 1 mM imidazole, 2 mM CaCl_2_, 1 mM DTT) for 1-2 hours at 4 °C. Beads were washed in 40 bead volumes calmodulin binding buffer and eluted in small fractions (300 µl) using calmodulin elution buffer (like calmodulin binding buffer, 2 mM CaCl_2_ is substituted with 20 mM EGTA). Single fractions were flash frozen in liquid nitrogen and stored at −80 °C until further usage in in vitro kinase assays.

### Yeast genetics (Strain construction, FRB system, pESCs, Integration/replacement constructs)

Yeast strains were constructed in the S288C or W303 (SCF strains) background. A list of all yeast strains used in this study can be found in **Table 4**, a list of all vectors used for generation of novel yeast strains can be found in **Table 3.** Yeast strain generation and methods were performed by standard procedures (Daniel et al., 2006). The anchor-away approach for characterization of Ame1 in SCF mutants or wild-type strain was performed as described (Haruki et al., 2008), using the ribosomal RPL13-FKBP12 anchor. Final rapamycin concentration in plates or liquid media was 1 μg/ml. Serial two-fold dilutions of overnight cultures were prepared on 96-well plates in minimal medium starting from OD_600_ of 0.4 for anchor-away approach or 0.5 for overexpression approach. The dilutions were spotted on YPD medium with and without rapamycin or on minimal medium with either S-glucose or S-raffinose + galactose (2 % each) and grown at 30 °C for 2-3 days. To confirm phenotypes observed in the serial dilution assays for Ame1 mutants, Ame1 hemizygous deletion strains were used to introduce Ame1-wild-type or phospho-mutants at an exogenous locus before haploid spores were produced. Pds1-13xMyc was integrated exogenously into haploid strains for cell cycle experiments. For overexpression of proteins, pESC two-micron plasmids were integrated in haploid wild-type or SCF mutant strains without integration into the genome and clone pools were used for further analyses. Selective pressure was used for maintenance of the plasmids. Expression of integrated proteins was checked for all created yeast strains by protein extraction from yeast (Kushnirov, 2000) and western blotting against the respective tags of individual proteins (see **Table 2**).

### GAL overexpression

For overexpression studies strains containing two-micron plasmids were grown overnight in YEPD. Next morning, cells were washed twice in YEP-raffinose (2 %) and incubated in YEP+R for 3 hours at 30 °C. Overexpression was induced with the addition of 2 % galactose and timepoints were taken after 0, 3 and 5 hours in YEP+RG. Protein extracts (Kushnirov, 2000) for western blotting analysis were prepared for each timepoint and accumulation of protein over time was visualized using anti-Flag M2 antibody for Ame1, anti c-myc (9E10) for Okp1 or Ctf19 and anti-HA for Mcm21.

### Cell cycle analysis (Synchronization, FACS)

For cell cycle experiments, cells were diluted in 80 ml of liquid YEPD with a starting OD_600_ = 0.2 and incubated for 1 hour at 30 °C. Next, cells were arrested in a G1-like state with alpha-factor for 2 hours at 30 °C. Afterwards, cells were washed extensively in YEPD + pronase (100 µg/ml) and in YEPD to remove the alpha-factor. Cells were than grown in fresh YEPD at 30 °C and protein extracts were prepared immediately and in intervals of 15 minutes using the method described in Kushnirov 2000. For FACS staining, cells for each timepoint were fixed in 95 % EtOH overnight at 4 °C. Fixed cells were washed in 50 mM Tris-HCl pH 7.5 and sonicated using an ultrasonic waterbath. RNA was digested using 0.2 mg/ml RNAse (**Sigma**) for 2 hours and 50 °C. Proteinase K (20 mg/ml pH 7.5; Roche) was added and cells were incubated further for 2-2.5 hours at 50 °C (Method described in (Haase and Reed, 2002)). Cell suspension was stained with 1 µM Sytox Green (Invitrogen) for 30 minutes at room temperature. Stained cells were stored at 4 °C and read out using the MaxQuant VJB machine.

### In vitro kinase reactions

For in vitro kinase reactions purified AO complexes (wild-type or different phospho-mutants in Ame1) were phosphorylated using Cdc28-Clb2. AO with a concentration of 5 µM was incubated with recombinant Cdc28-Clb2, 1 µl 1 mM cold ATP, 1 µl radioactive labeled ATP (10 µCi/µl) and 4x kinase buffer (80 mM Hepes pH 7,5, 400 mM KCl, 40 mM MgCl_2_, 40 mM MnCl_2_, 100 mM β-Glycerolphosphate, 4 mM DTT) in a final volume of 20 µl. The reaction was incubated for 30 minutes at 30 °C and stopped by adding 6x SDS loading dye. The whole reaction was loaded on a SDS-PAGE and stained with Coomassie Brilliant Blue. After destaining the gel overnight, an X-ray film was exposed for 5-20 minutes and developed with CAWOMAT 200 IR machine.

For kinase assays with Ame1-Okp1 complexes and Mtw1c or MN, recombinant proteins were preincubated in a molar ratio of 2:1, where always 10 µM of AO and 5 µM Mtw1c or MN were used. Mixed protein samples were incubated on ice for 30 minutes, followed by adding all other components described above and starting the kinase reaction.

For the quantitative phosphorylation analysis presented in **Supplementary Figure 1**, 1 µM AO complex was phosphorylated with 0.3 nM cyclin-Cdc28 complex in a buffer containing 50 mM Hepes-KOH, pH 7.4, 150 mM NaCl, 5 mM MgCl_2_, 20 mM imidazole, 2% glycerol, 0.2 mg/ml BSA, 500 nM Cks1, and 500 µM ATP [(with added [γ-^32^P]-ATP (Hartmann Analytic)]. The reaction was stopped at 10 minutes (initial velocity condition, less than 10% of initial substrate turnover) by addition of SDS loading dye. The samples loaded on SDS-PAGE and γ-^32^P phosphorylation signals were detected using Amersham Typhoon 5 Biomolecular Imager (GE Healthcare Life Sciences).

### Computational details of molecular dynamics simulations

The crystal structure of Cdc4 with a phosphopeptide (PDB ID: 1NEX, (Orlicky et al., 2003) resolution: 2.7 Å) was used for the initial coordinates of our models. The reported complex comprises Cdc4, Skp1, and a linker region. For the present study, we removed Skp1 and the linker region as both are not directly involved in peptide binding. Further, the ions and crystallization water molecules were removed. The selenomethionine modifications in the protein structure were replaced by the parent methionine residues.

The peptides under study were modeled using the PEP-FOLD3.5 webserver (Lamiable et al., 2016). For performing the simulations, the control peptide in the crystal structure was removed and manually replaced by the candidate peptides, with further optimization using the FlexPepDock webserver (London et al., 2011). All the simulations were performed using the AMBER software suite (Case et al., 2014). The protein counterpart and the regular amino acids in the peptide were simulated with the modified ff99SB AMBER force field (Maier et al., 2015). The phosphorylated serine and threonine parameters were taken from the literature (Homeyer et al., 2006). Gaussian accelerated molecular dynamics (GaMD) simulations (Miao et al., 2015) were performed in explicit solvent using a TIP3P water (Jorgensen et al., 1983) box with sufficient counterions (Na^+^/Cl^-^) to ensure the overall electroneutrality of the system. The parameters of the ions were taken from the literature (Joung and Cheatham, 2008). Periodic boundary conditions were implemented along with Particle Mesh Ewald (PME) for computing the long-range electrostatic interactions (Cheatham et al., 1995). The systems were minimized in two steps (using 10,000 conjugate gradient and 10,000 steepest descent cycles for both steps). In step1, the protein-peptide complex was restrained using a force constant of 50 kcal/mol A^−2^, and only the ions and solvent molecules were allowed to relax. In step 2, the restraints were removed, and the whole assembly was allowed to relax. The minimized systems were heated to room temperature (40 ps) and underwent equilibration (5 ns). Finally, GaMD simulations were performed for 100 ns (3 replicas, NPT). The trajectories were processed using the Cpptraj code (Roe and Cheatham, 2013), and the downloadable version of the Ligplot tool was used to create 2D interaction plots (Wallace et al., 1995). For the clustering of the GaMD replicas, we utilized the concepts of hierarchical agglomerative clustering (Shao et al., 2007), with a cut-off distance of 3 Å between any identified clusters.

## 2 Antibodies used in western blotting

2: Antibodies used in western blotting
| Antibody | Provider | Dilution |
| --- | --- | --- |
| Anti-FLAG M2 Monoclonal<br>Antibody-Peroxidase<br>Conjugate | Sigma | 1:10000 |
| Phosphoglycerate Kinase<br>Monoclonal Antibody | Invitrogen | 1:10000 |
| c-myc (9E10) monoclonal | BioLegend | 1:1000 |
| Peroxidase-conjugated<br>AffiniPure Goat Anti-Mouse<br>IgG | JacksonImmunoResearch | 1:5000 |
| Anti-Tubulin-HRP | Santa Cruz Biotechnologies | 1:1000 |
| Anti-HA | Covance | 1:1000 or 1:2000 |

## 3 Vectors for protein expression and yeast strain generation

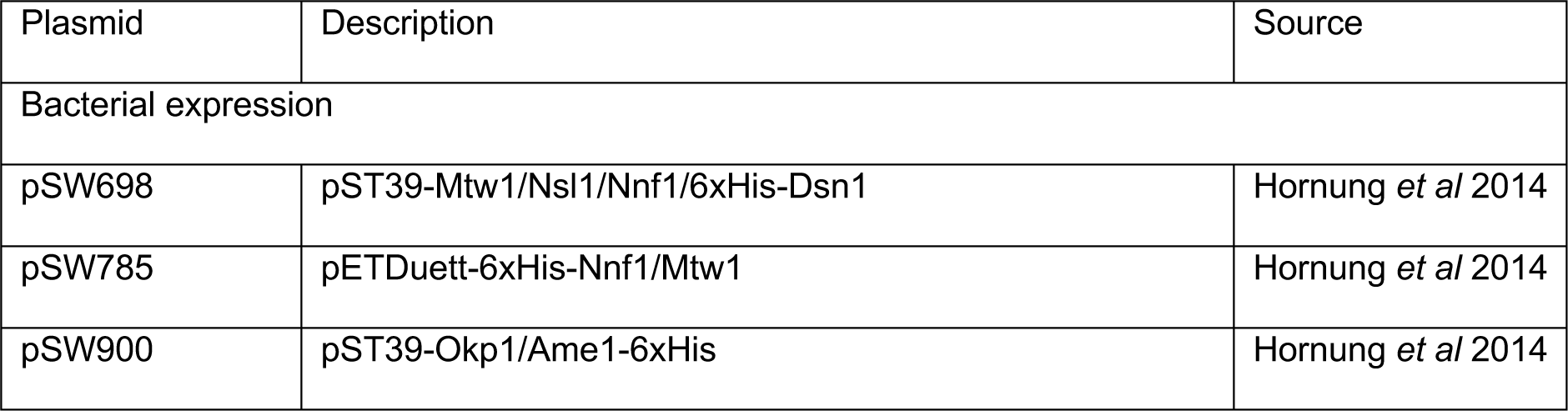
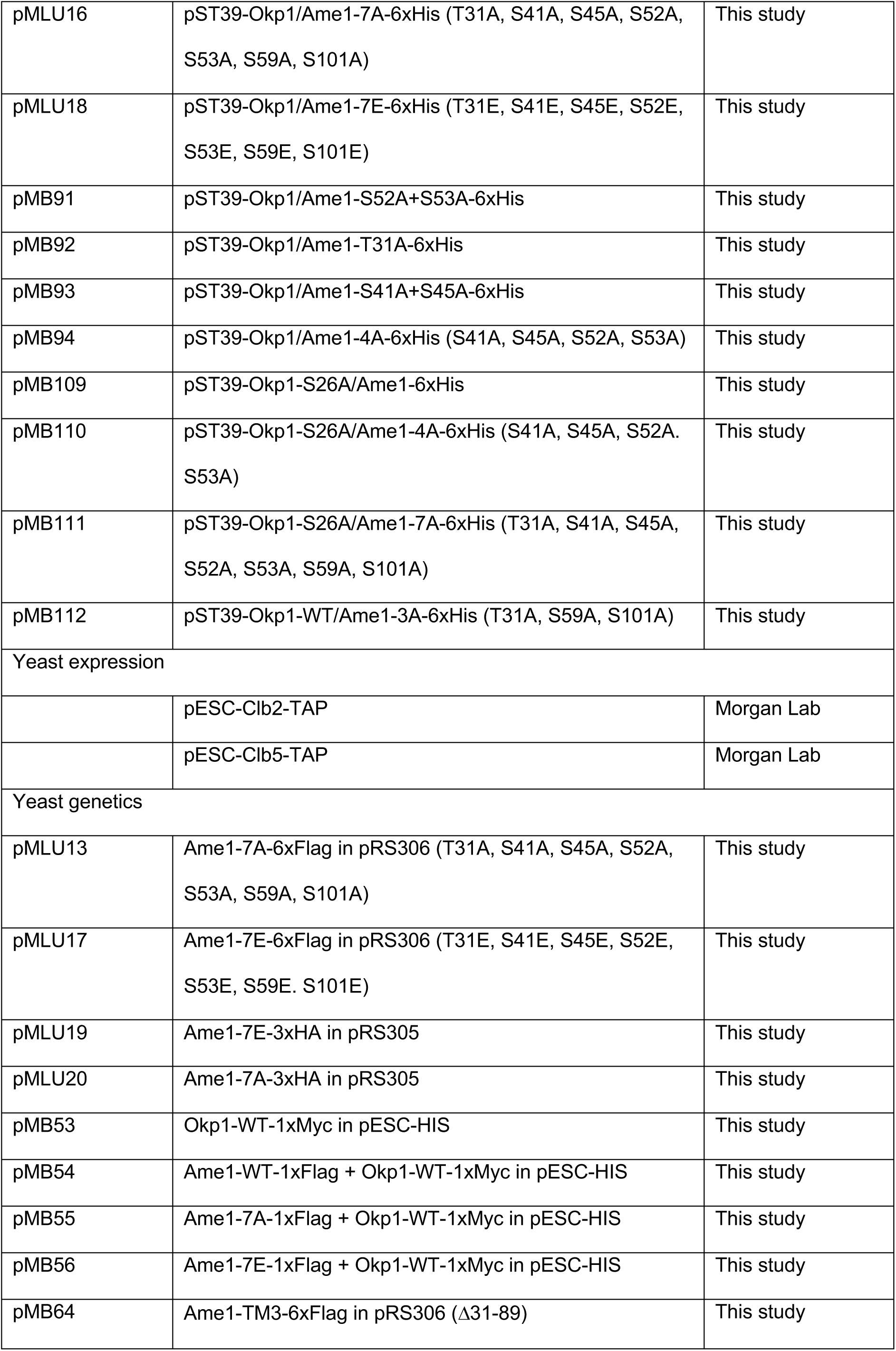
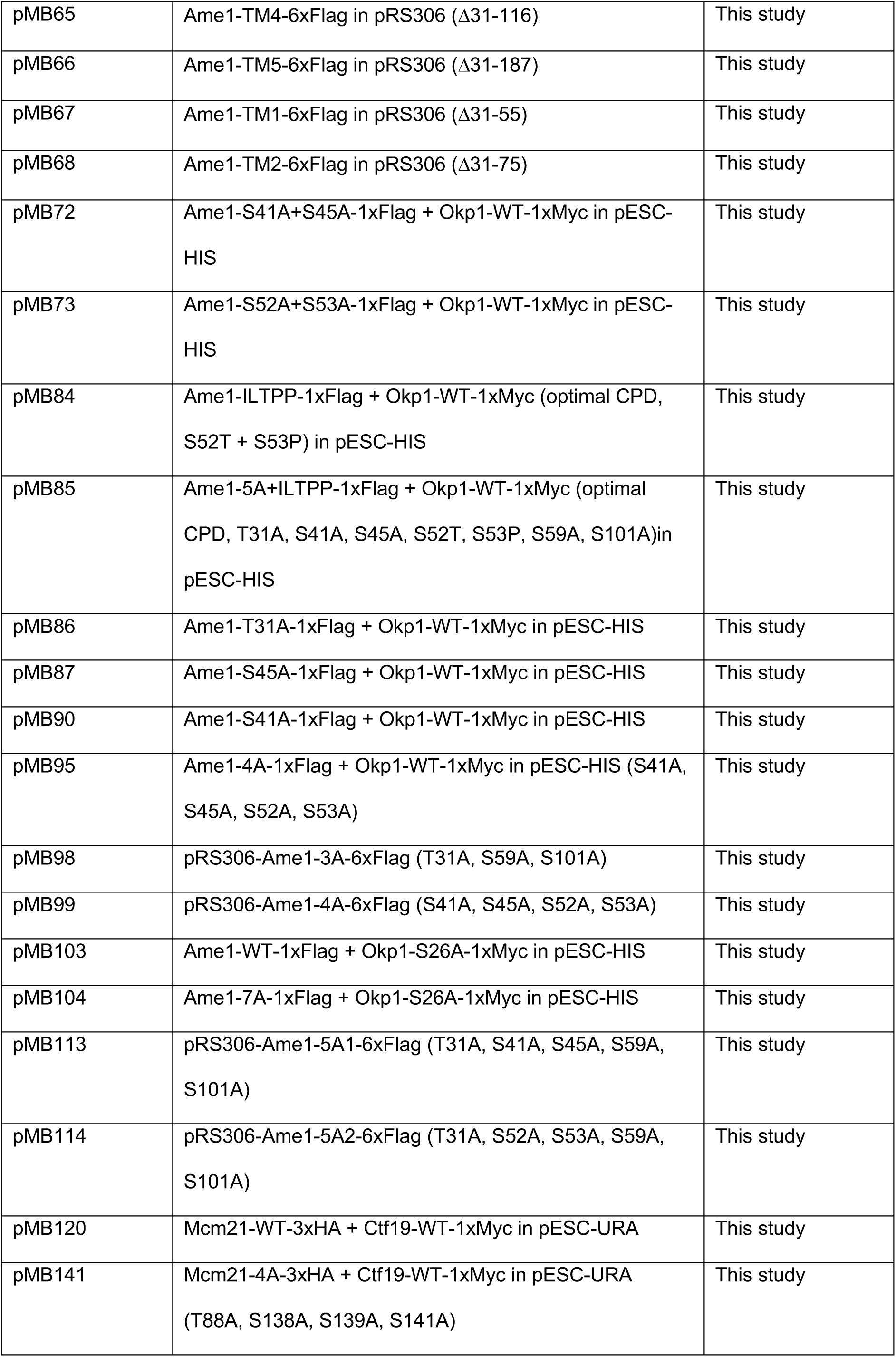
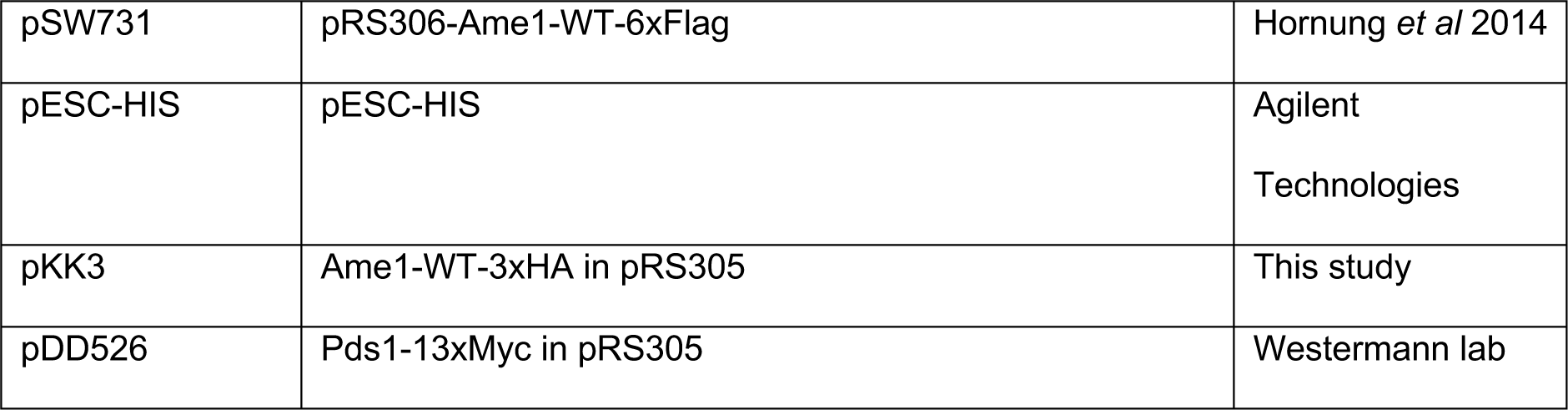

3: Vectors for protein expression and yeast strain generation
| Plasmid | Description | Source |
| --- | --- | --- |
| Bacterial expression |  |  |
| pSW698 | pST39-Mtw1/Nsl1/Nnf1/6xHis-Dsn1 | Hornung *et al* 2014 |
| pSW785 | pETDuett-6xHis-Nnf1/Mtw1 | Hornung *et al* 2014 |
| pSW900 | pST39-Okp1/Ame1-6xHis | Hornung *et al* 2014 |
| pMLU16 | pST39-Okp1/Ame1-7A-6xHis (T31A, S41A, S45A, S52A, S53A, S59A, S101A) | This study |
| pMLU18 | pST39-Okp1/Ame1-7E-6xHis (T31E, S41E, S45E, S52E, S53E, S59E, S101E) | This study |
| pMB91 | pST39-Okp1/Ame1-S52A+S53A-6xHis | This study |
| pMB92 | pST39-Okp1/Ame1-T31A-6xHis | This study |
| pMB93 | pST39-Okp1/Ame1-S41A+S45A-6xHis | This study |
| pMB94 | pST39-Okp1/Ame1-4A-6xHis (S41A, S45A, S52A, S53A) | This study |
| pMB109 | pST39-Okp1-S26A/Ame1-6xHis | This study |
| pMB110 | pST39-Okp1-S26A/Ame1-4A-6xHis (S41A, S45A, S52A, S53A) | This study |
| pMB111 | pST39-Okp1-S26A/Ame1-7A-6xHis (T31A, S41A, S45A, S52A, S53A, S59A, S101A) | This study |
| pMB112 | pST39-Okp1-WT/Ame1-3A-6xHis (T31A, S59A, S101A) | This study |
| Yeast expression |  |  |
|  | pESC-Clb2-TAP | Morgan Lab |
|  | pESC-Clb5-TAP | Morgan Lab |
| Yeast genetics |  |  |
| pMLU13 | Ame1-7A-6xFlag in pRS306 (T31A, S41A, S45A, S52A, S53A, S59A, S101A) | This study |
| pMLU17 | Ame1-7E-6xFlag in pRS306 (T31E, S41E, S45E, S52E, S53E, S59E, S101E) | This study |
| pMLU19 | Ame1-7E-3xHA in pRS305 | This study |
| pMLU20 | Ame1-7A-3xHA in pRS305 | This study |
| pMB53 | Okp1-WT-1xMyc in pESC-HIS | This study |
| pMB54 | Ame1-WT-1xFlag + Okp1-WT-1xMyc in pESC-HIS | This study |
| pMB55 | Ame1-7A-1xFlag + Okp1-WT-1xMyc in pESC-HIS | This study |
| pMB56 | Ame1-7E-1xFlag + Okp1-WT-1xMyc in pESC-HIS | This study |
| pMB64 | Ame1-TM3-6xFlag in pRS306 (Δ31-89) | This study |
| pMB65 | Ame1-TM4-6xFlag in pRS306 (Δ31-116) | This study |
| pMB66 | Ame1-TM5-6xFlag in pRS306 (Δ31-187) | This study |
| pMB67 | Ame1-TM1-6xFlag in pRS306 (Δ31-55) | This study |
| pMB68 | Ame1-TM2-6xFlag in pRS306 (Δ31-75) | This study |
| pMB72 | Ame1-S41A+S45A-1xFlag + Okp1-WT-1xMyc in pESC-HIS | This study |
| pMB73 | Ame1-S52A+S53A-1xFlag + Okp1-WT-1xMyc in pESC-HIS | This study |
| pMB84 | Ame1-ILTPP-1xFlag + Okp1-WT-1xMyc (optimal CPD, S52T + S53P) in pESC-HIS | This study |
| pMB85 | Ame1-5A+ILTPP-1xFlag + Okp1-WT-1xMyc (optimal CPD, T31A, S41A, S45A, S52T, S53P, S59A, S101A)in pESC-HIS | This study |
| pMB86 | Ame1-T31A-1xFlag + Okp1-WT-1xMyc in pESC-HIS | This study |
| pMB87 | Ame1-S45A-1xFlag + Okp1-WT-1xMyc in pESC-HIS | This study |
| pMB90 | Ame1-S41A-1xFlag + Okp1-WT-1xMyc in pESC-HIS | This study |
| pMB95 | Ame1-4A-1xFlag + Okp1-WT-1xMyc in pESC-HIS (S41A, S45A, S52A, S53A) | This study |
| pMB98 | pRS306-Ame1-3A-6xFlag (T31A, S59A, S101A) | This study |
| pMB99 | pRS306-Ame1-4A-6xFlag (S41A, S45A, S52A, S53A) | This study |
| pMB103 | Ame1-WT-1xFlag + Okp1-S26A-1xMyc in pESC-HIS | This study |
| pMB104 | Ame1-7A-1xFlag + Okp1-S26A-1xMyc in pESC-HIS | This study |
| pMB113 | pRS306-Ame1-5A1-6xFlag (T31A, S41A, S45A, S59A, S101A) | This study |
| pMB114 | pRS306-Ame1-5A2-6xFlag (T31A, S52A, S53A, S59A, S101A) | This study |
| pMB120 | Mcm21-WT-3xHA + Ctf19-WT-1xMyc in pESC-URA | This study |
| pMB141 | Mcm21-4A-3xHA + Ctf19-WT-1xMyc in pESC-URA (T88A, S138A, S139A, S141A) | This study |
| pSW731 | pRS306-Ame1-WT-6xFlag | Hornung *et al* 2014 |
| pESC-HIS | pESC-HIS | Agilent Technologies |
| pKK3 | Ame1-WT-3xHA in pRS305 | This study |
| pDD526 | Pds1-13xMyc in pRS305 | Westermann lab |

## 3 Yeast strains

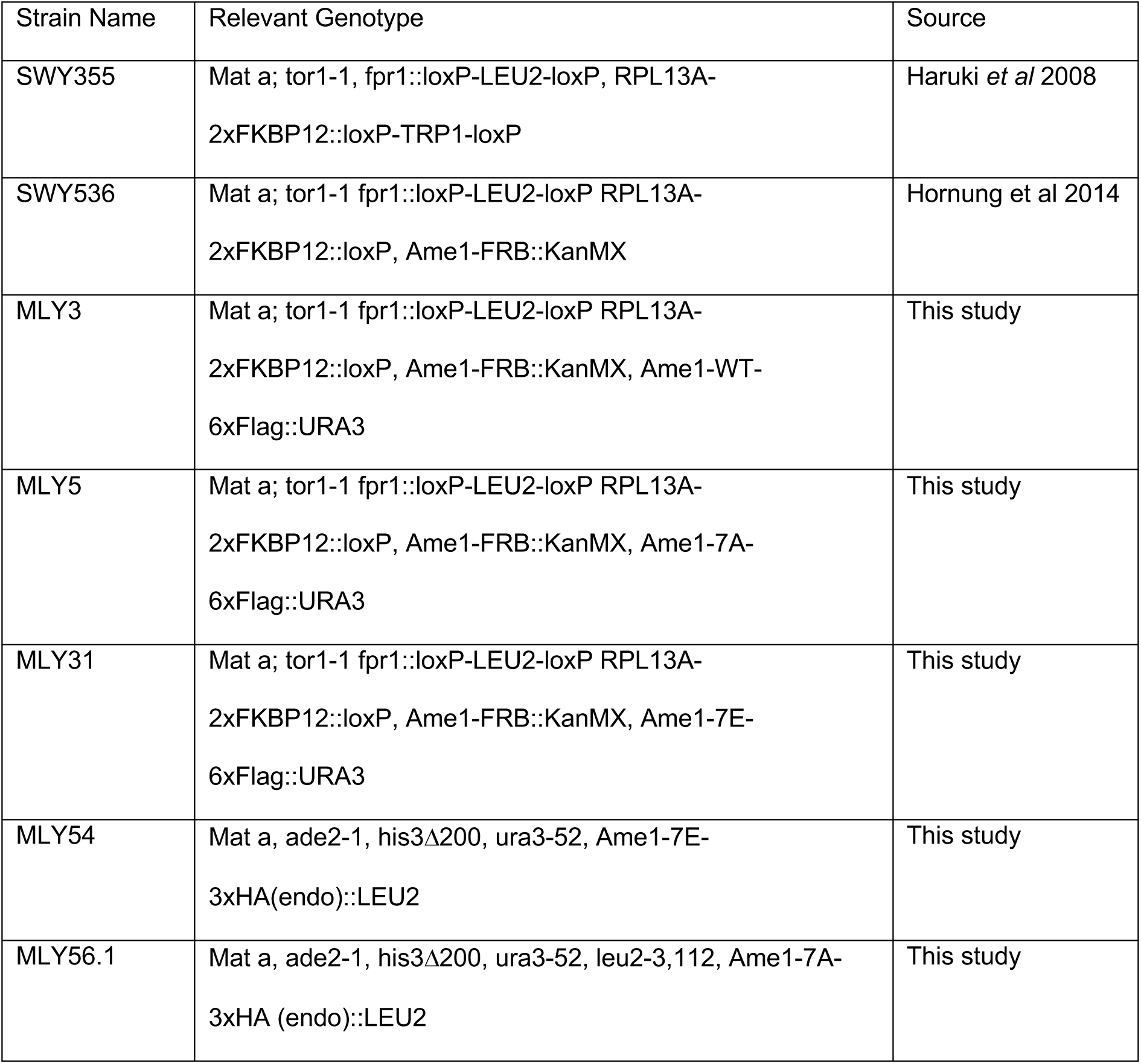
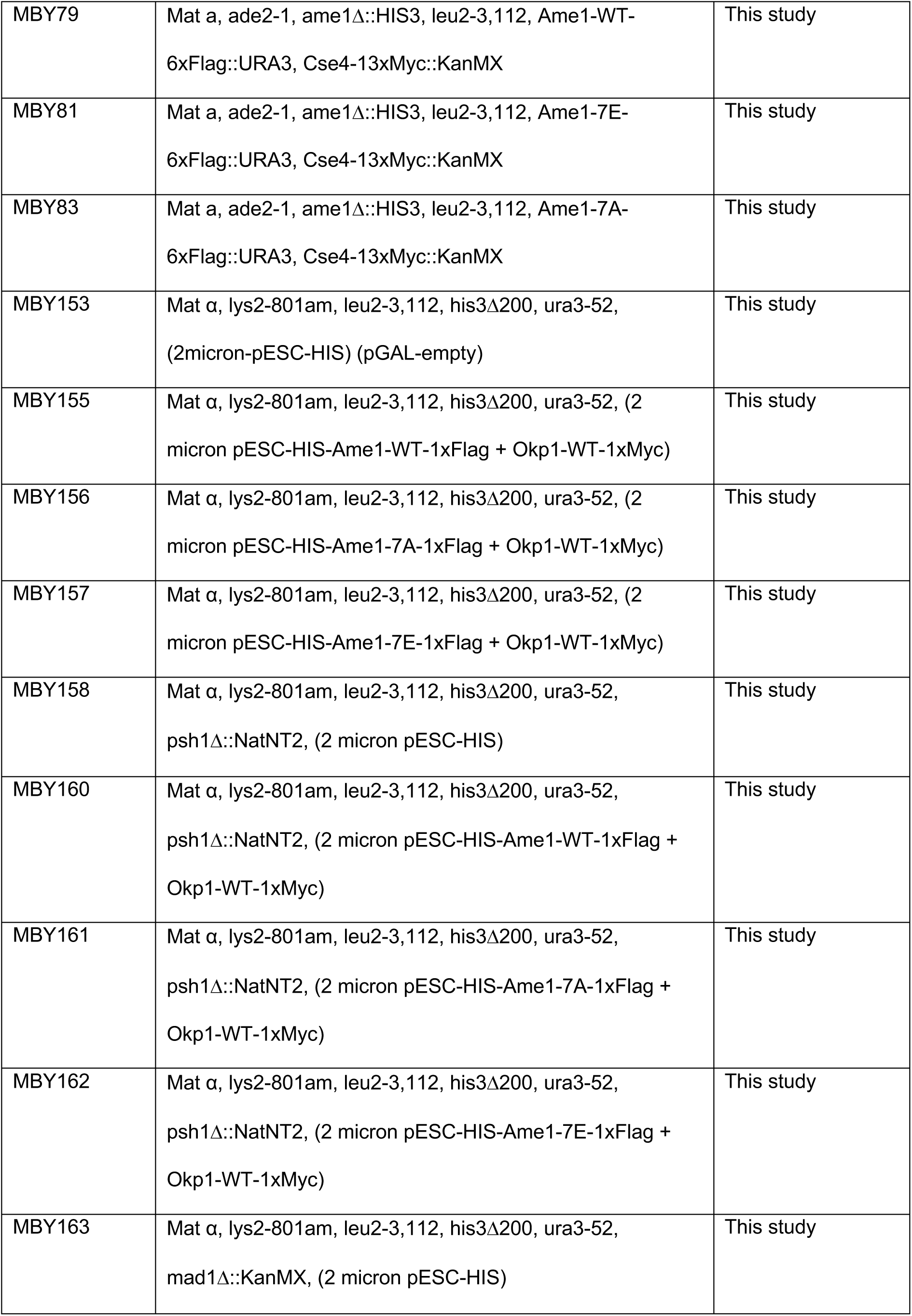
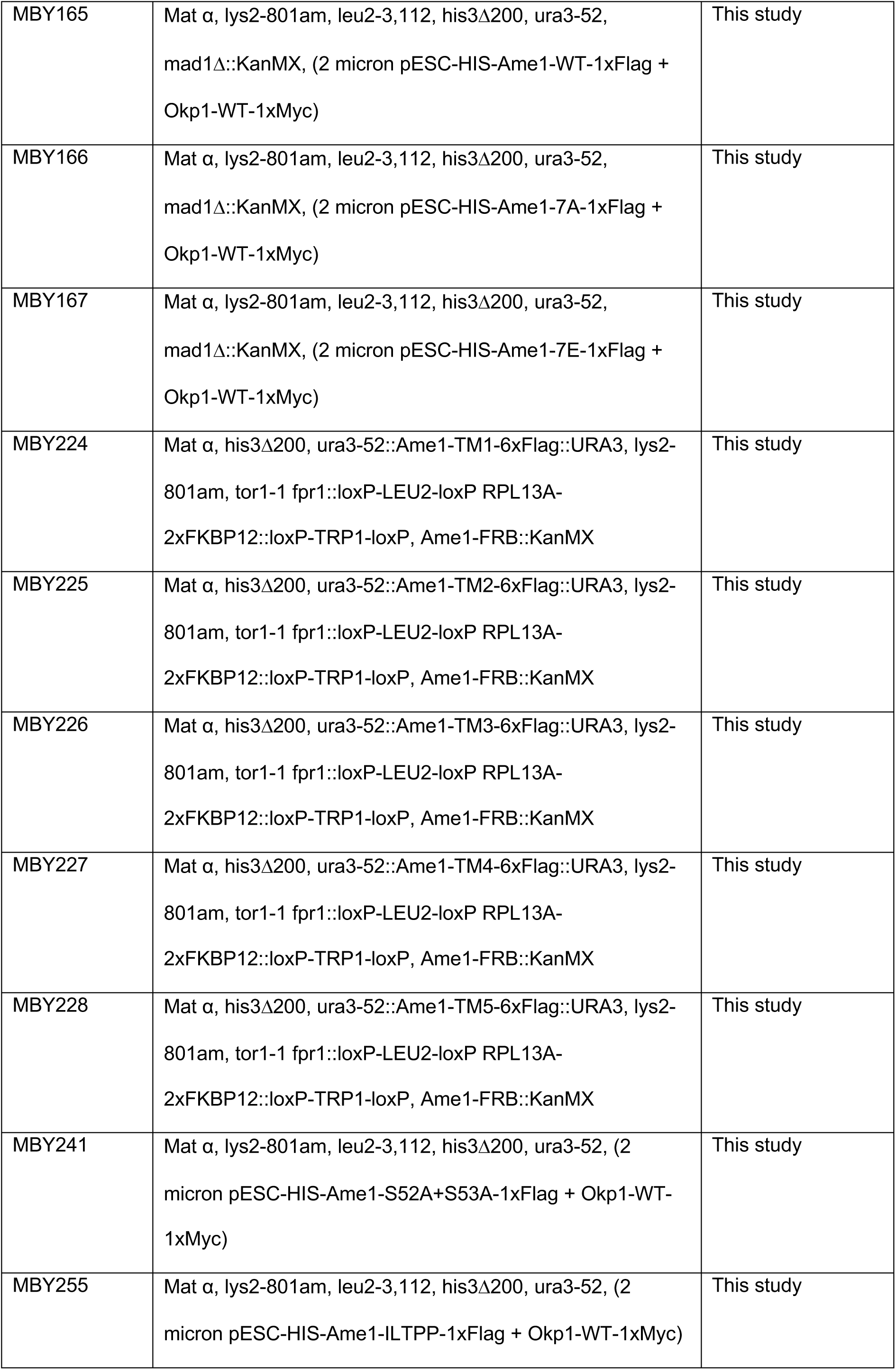
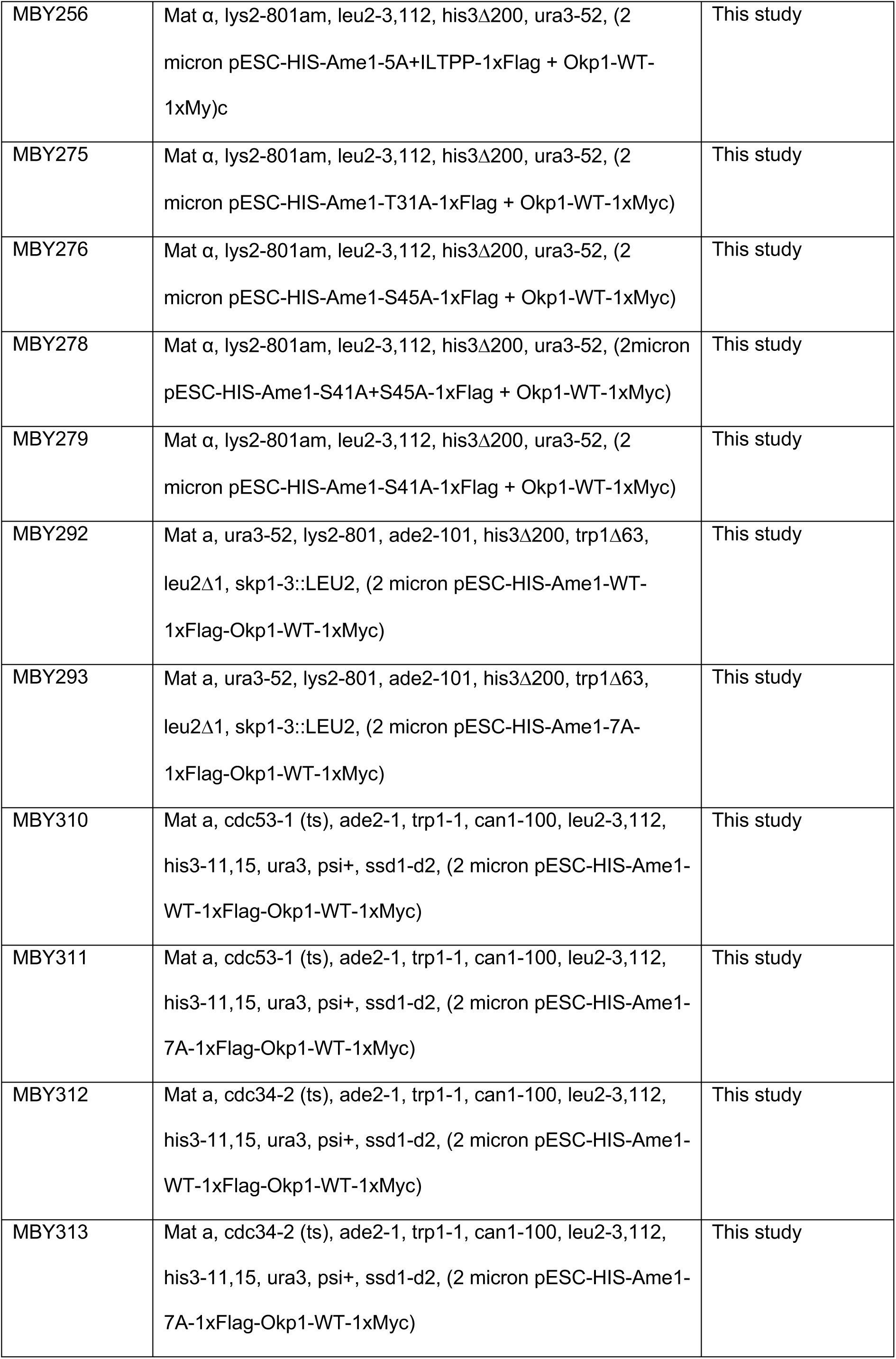
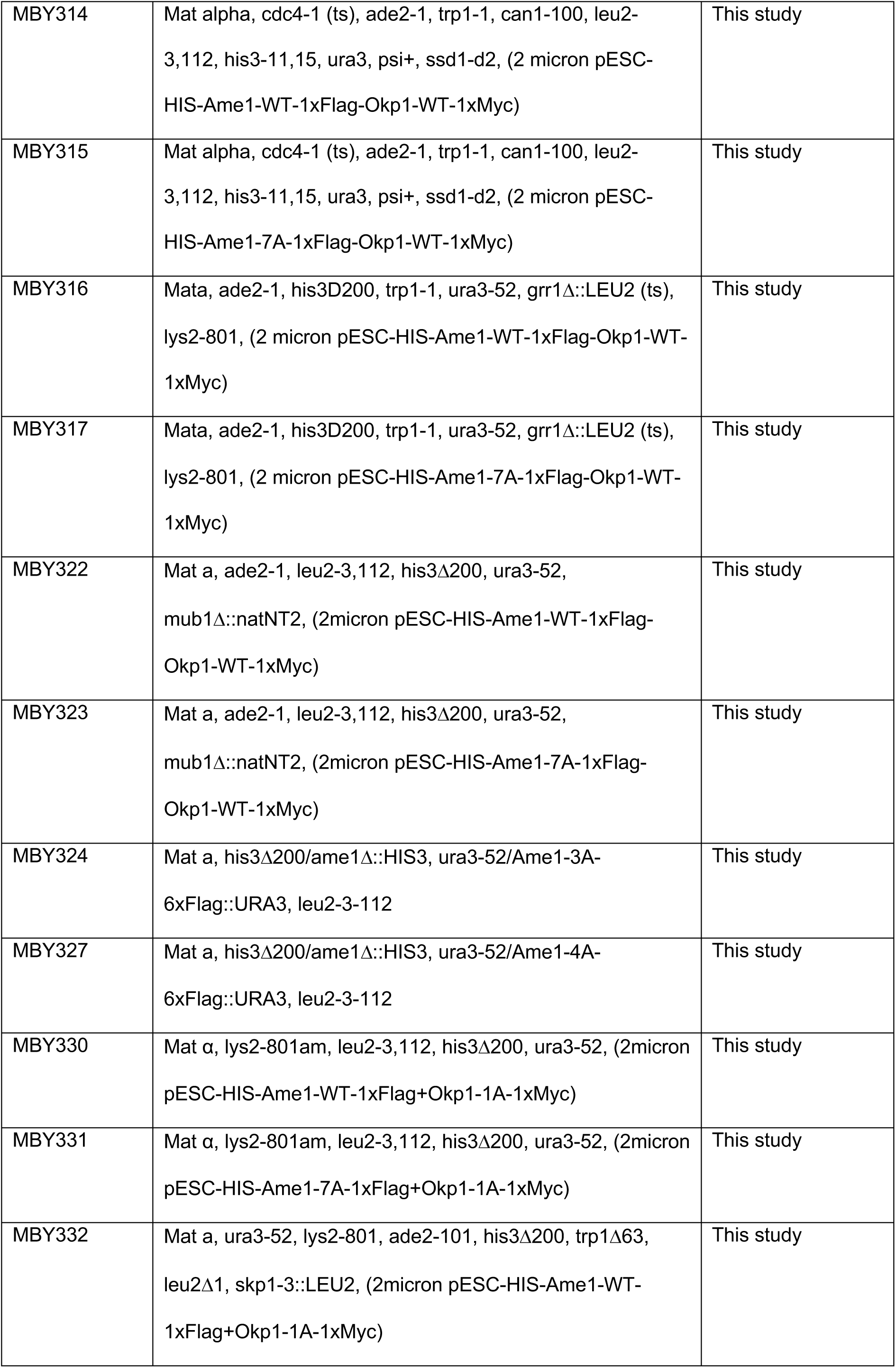
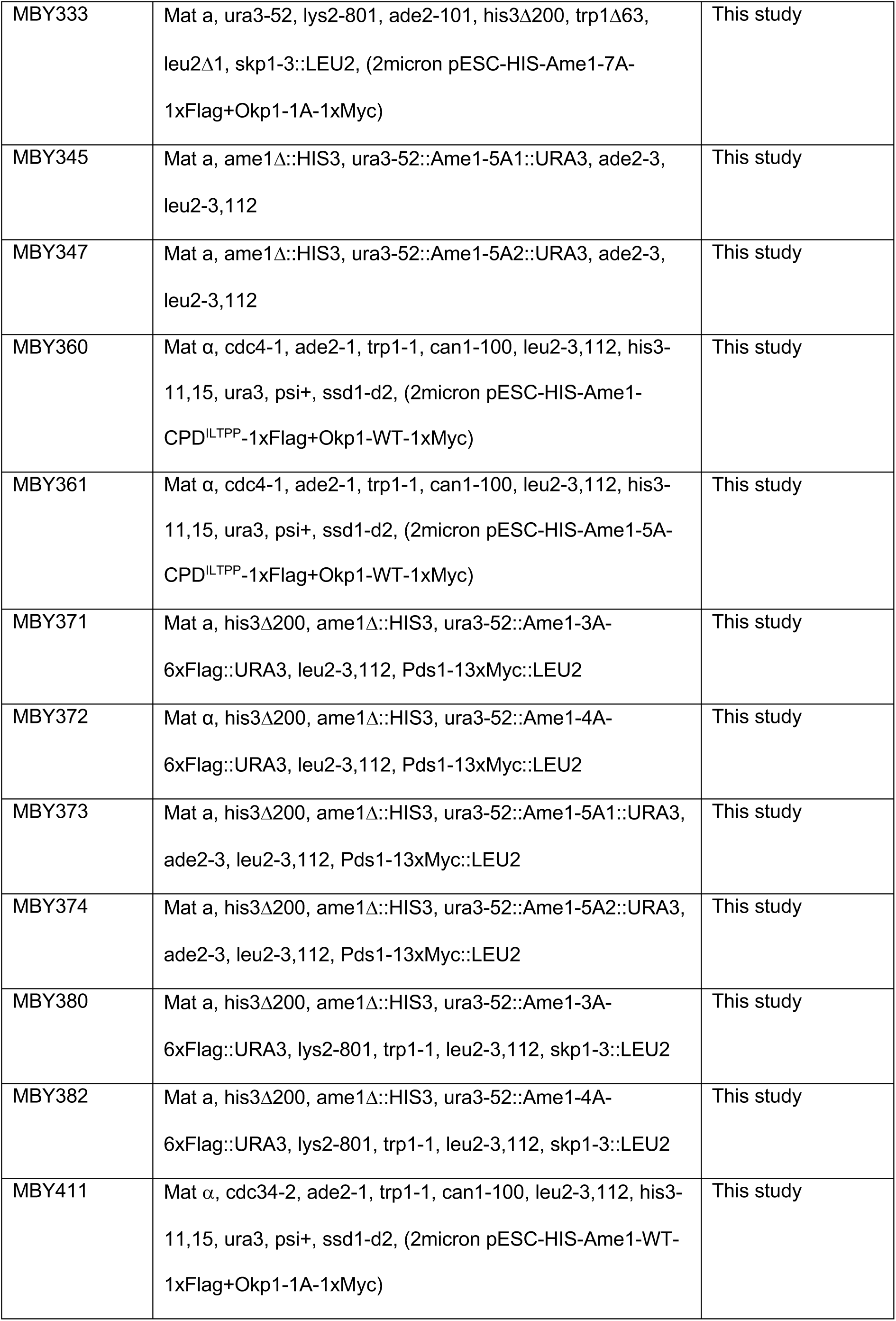
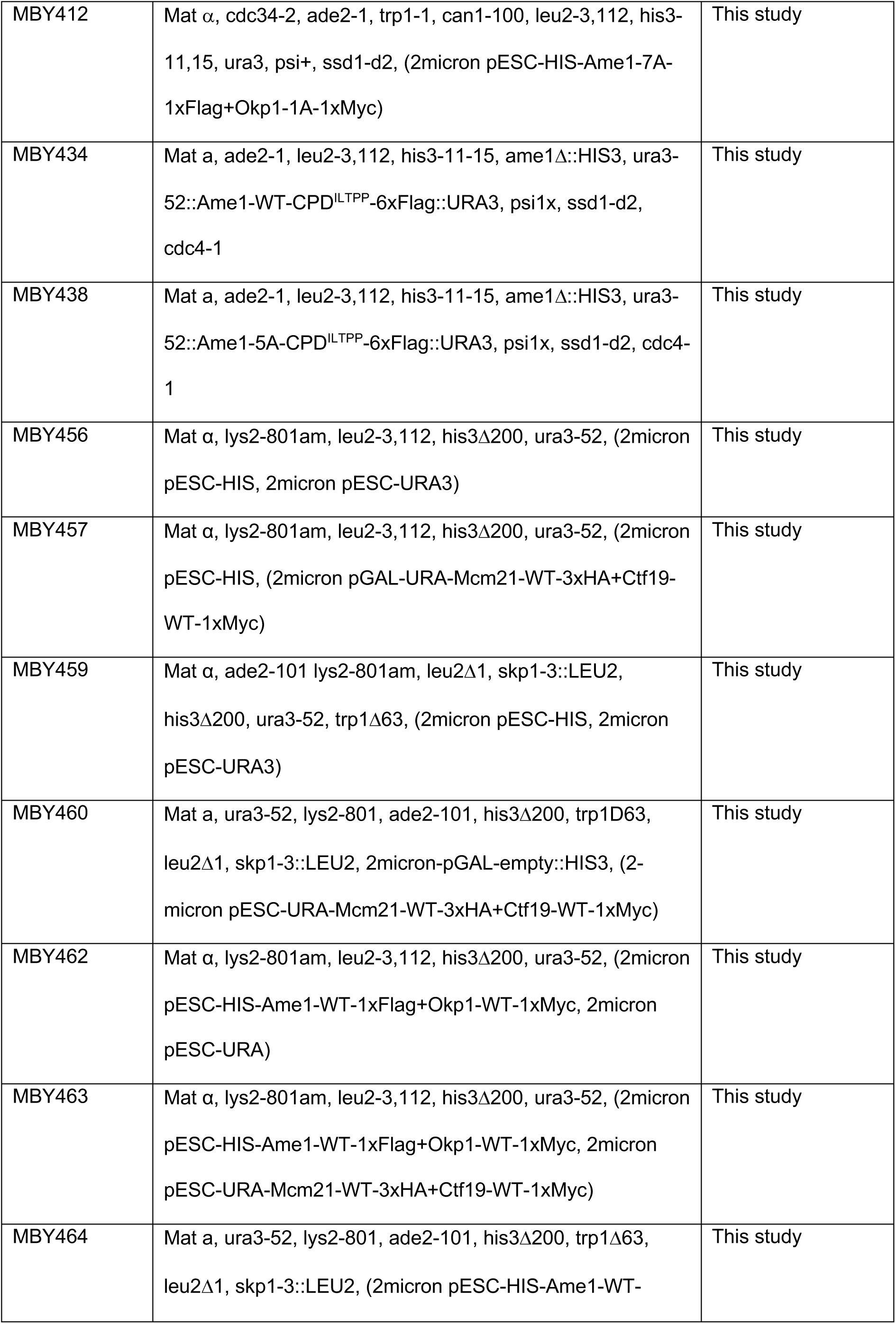
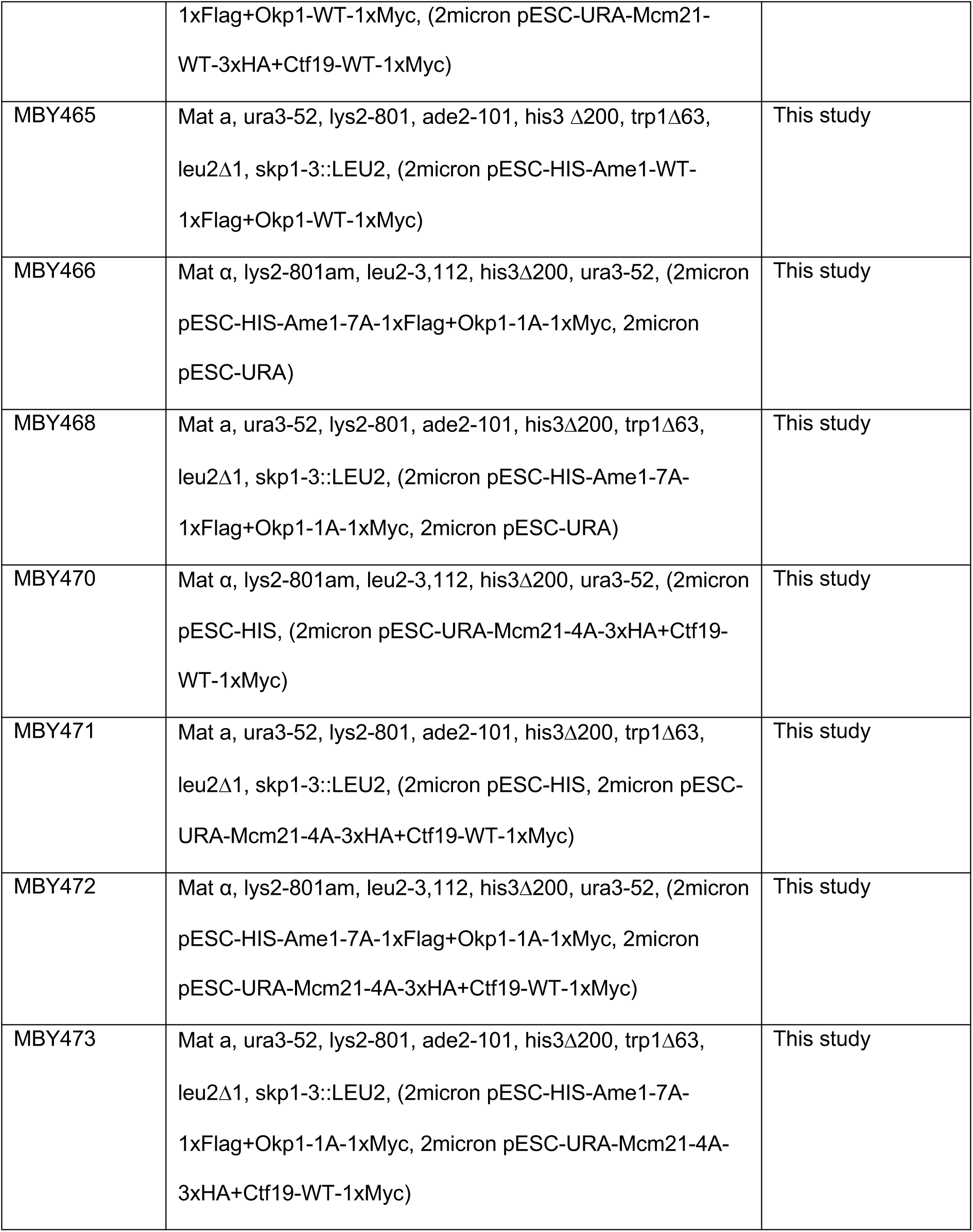

## Acknowledgments

The authors wish to thank Mathias Peter and Sue Biggins for providing strains and plasmids. We thank Maren Soldierer and Tabea Graszynski for initial experimental contributions and all members of the Westermann lab for discussions. This work received support from the German Research Foundation (DFG), grant no. WE-2886/2. E. S.-G. was also supported by Germany’s Excellence Strategy – EXC 2033 – 390677874 – RESOLV, funded by the DFG. S.W. and E. S.-G. received funding from the collaborative research center CRC 1093 “Supramolecular Chemistry on Proteins” (Subprojects A8 and B7), funded by the DFG. E. S.-G. acknowledges the computational time provided by the supercomputer magnitUDE of the University of Duisburg-Essen.

## Author contributions

MB performed biochemical and genetic experiments, supported by KK, AD, KJ, SH. KM performed and analyzed mass spectrometry experiments. PP performed the GaMD simulations, analyzed the results and wrote the computational section together with ESG. ESG designed the computational work, analyzed the simulations, and wrote the computational section together with PP. MÖ performed quantitative analysis of Cyclin-CDK phosphorylation, supervised by ML. MB and SW designed research, analyzed data and wrote the manuscript with input from all authors.

## Supplementary Figures

**Supplementary Figure 1.**
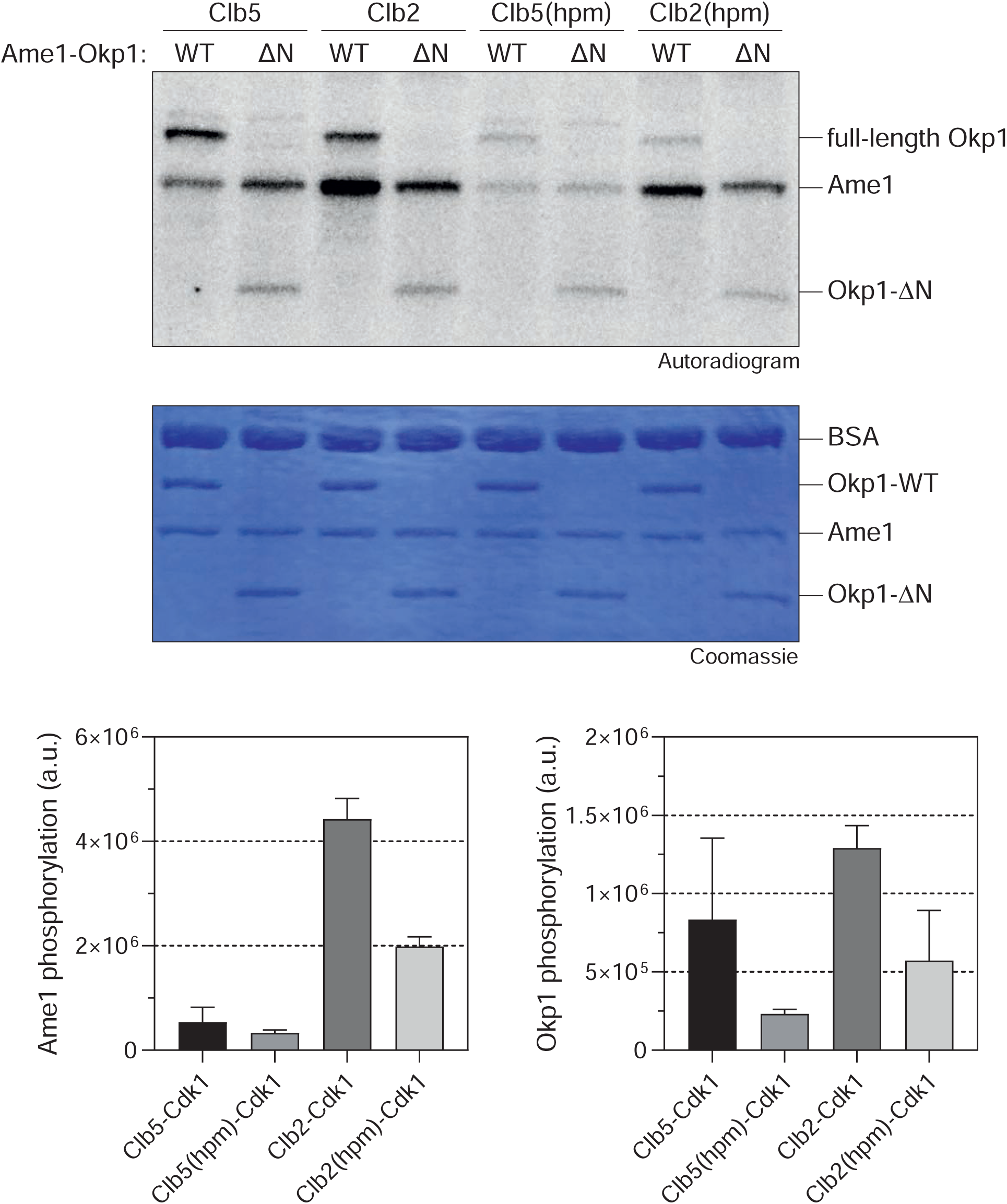
Quantitative phosphorylation analysis of recombinant Ame1-Okp1 by S-Cdk1 and M-Cdk1 complexes. In vitro phosphorylation of recombinant AO with Cdc28-Clb5 and Cdc28-Clb2 complexes or their hydrophobic patch mutants (hpm) which eliminate docking dependent phosphophorylation was analyzed. Autoradiogram of the kinase reaction is shown at the top, corresponding Coomassie stained gel is shown below. hpm: hydrophobic patch mutant. ΔN: N-terminal truncation mutant of Okp1. Quantification of phosphorylation for Ame1 and Okp1 is given below.

**Supplementary Figure 2.**
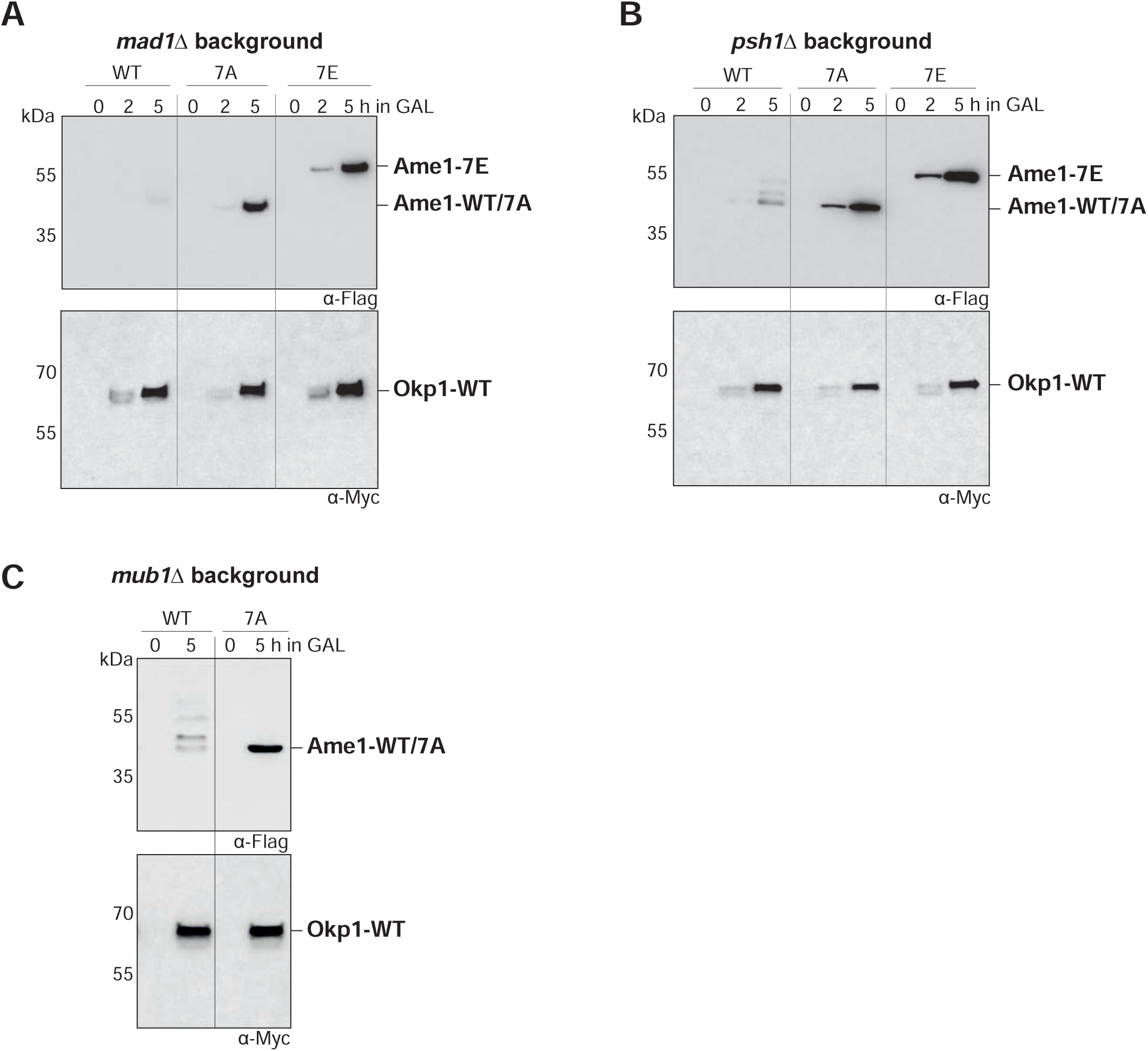
**A.** Western blot analysis of overexpression of Ame1-WT and phosphorylation mutants from a Galactose-inducible promoter in a *psh1Δ* background strain. Note that Ame1 levels remain low in this background. Ame1 phosphorylation mutants show the same accumulation effect as seen in wild-type background. **B**. Same experiment as **A** in a checkpoint deficient *mad1Δ* strain background. **C:** Overexpression of Ame1-WT and phosphorylation mutants in a deletion mutant of *mub1*, which is one subunit of the Ubr2/Mub1 ubiquitin ligase complex implicated in the regulation of Dsn1.

**Supplementary Figure 3.**
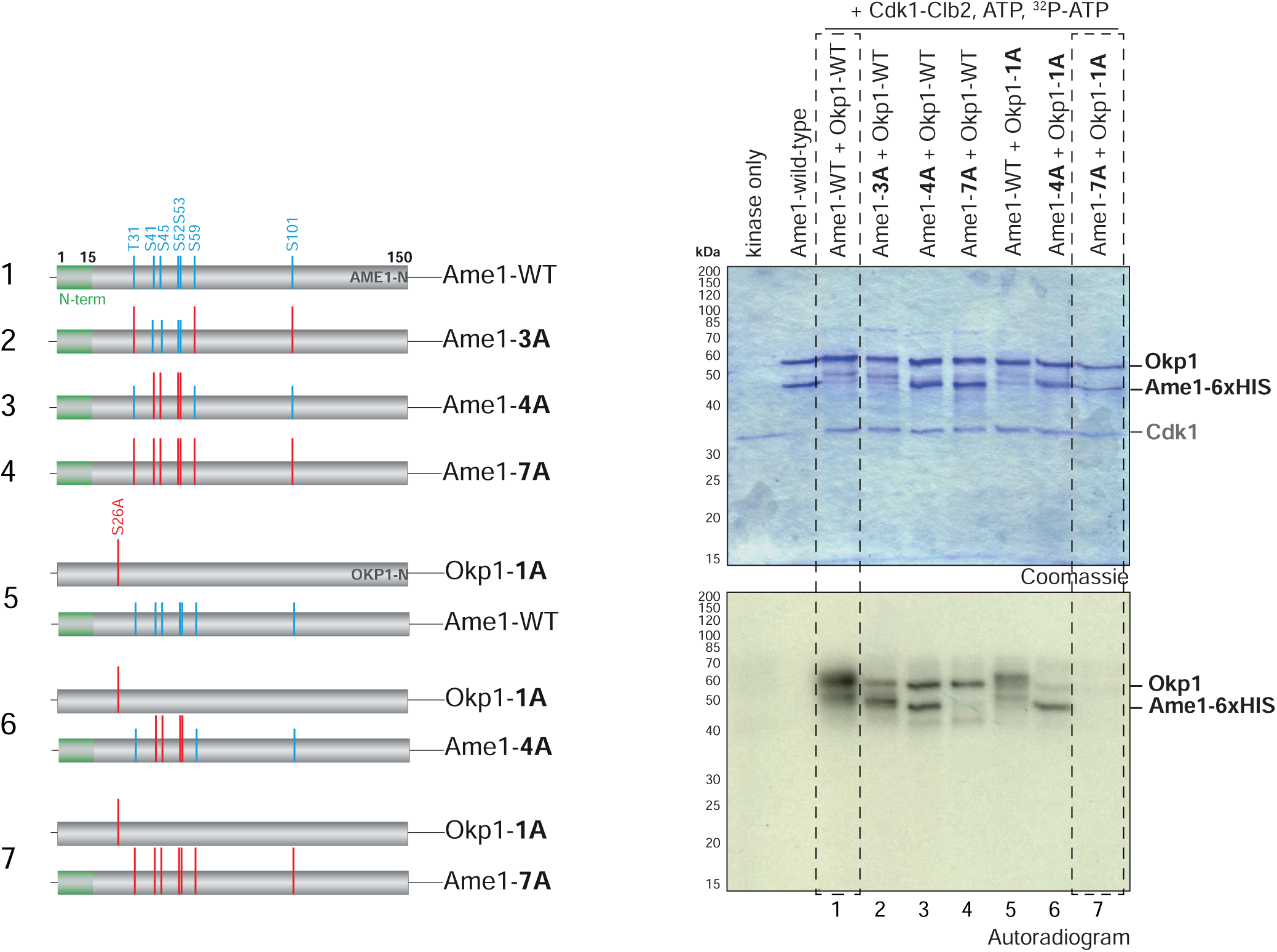
Additional analysis of AO in vitro phosphorylation by Cdc28-Clb2. Left panel: Scheme of the used recombinant AO complexes. For the Okp1-1A mutant, the Cdk1 site Ser26 was mutated to alanine. Right panel: Coomassie-stained gel and corresponding autoradiography of in vitro kinase assays with the indicated AO variants. Note that Ame1-7A in combination with Okp1-1A completely eliminates phosphorylation.

**Supplementary Figure 4:**
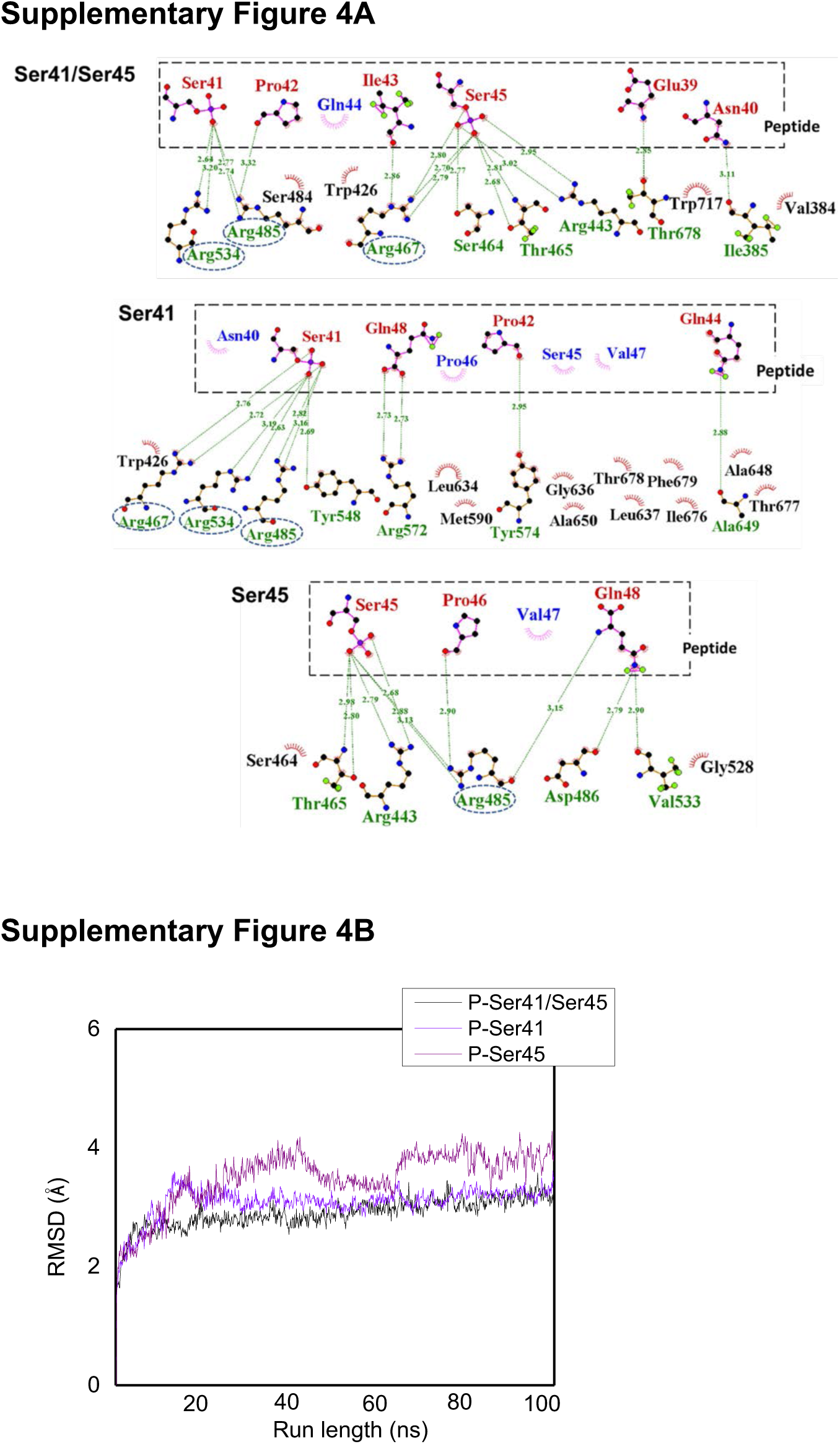
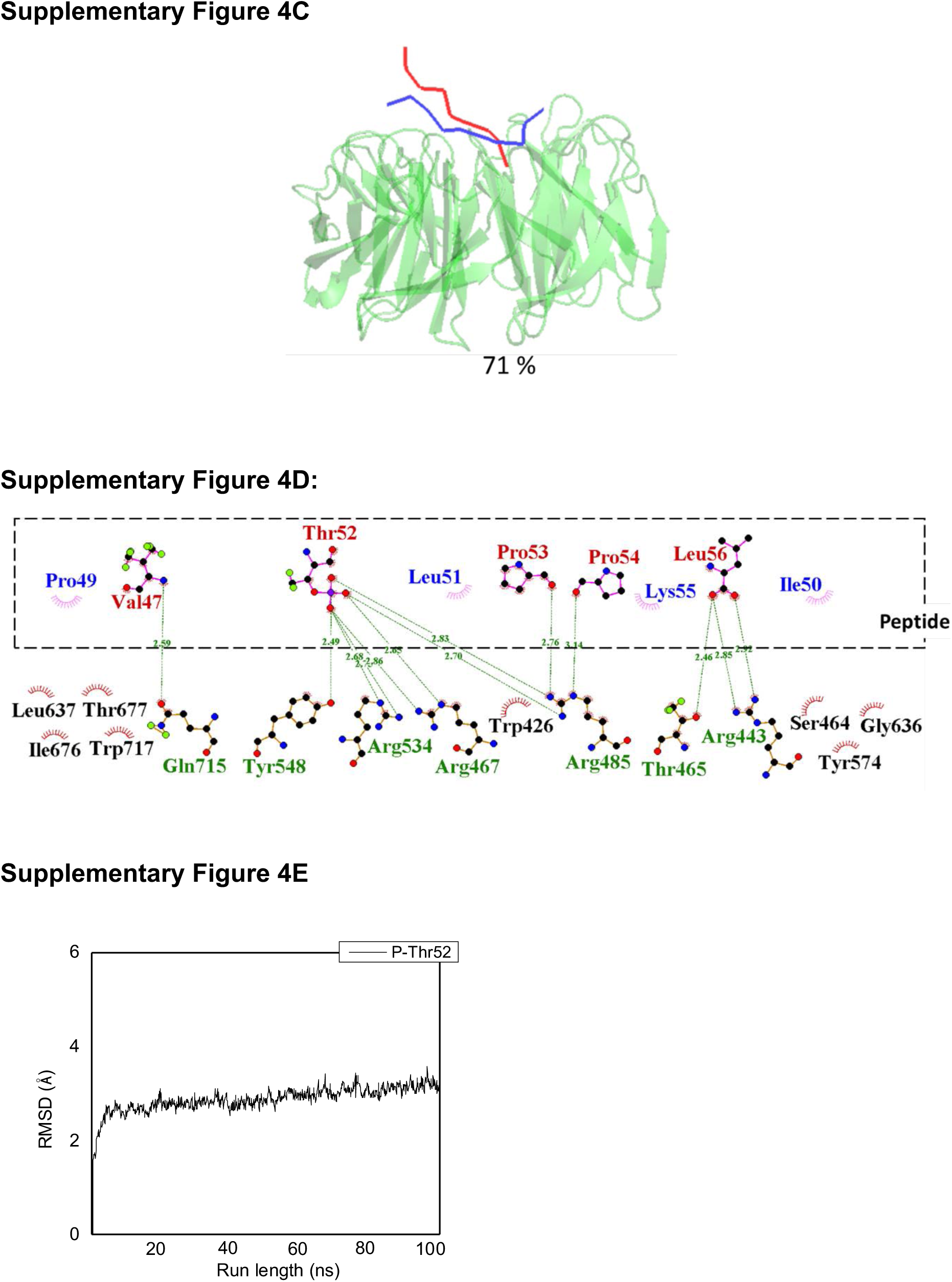
Additional analysis of peptide-Cdc4 interactions by Gaussian accelerated MD simulations. **A:** Interactions of the doubly phosphorylated Ame1 peptide Ser41/Ser45, as well as the monophosphorylated Ser41 and Ser45 peptides with the Cdc4 WD40 domain are shown. **B:** RMSD fluctuations for the doubly phosphorylated Ser41/Ser45 (P-Ser41/Ser45), monophosphorylated Ser41 (P-Ser41) and Ser45 (P-Ser45) complexed with Cdc4. **C:** The representative frame from the dominating cluster of Ame1-CPD^ILTPP^ – Cdc4 complex simulations (Peptide: VQPIL**T**PPKL, shown in blue) is aligned with the crystal structure taken for the study (the peptide counterpart is shown in red ribbon). The population of the dominating cluster is also shown. **D:** Interactions of phosphorylated threonine peptide with the Cdc4 protein. **E:** RMSD fluctuations displayed by Ame1-CPD^ILTPP^ complexed with Cdc4.

**Supplementary Figure 5.**
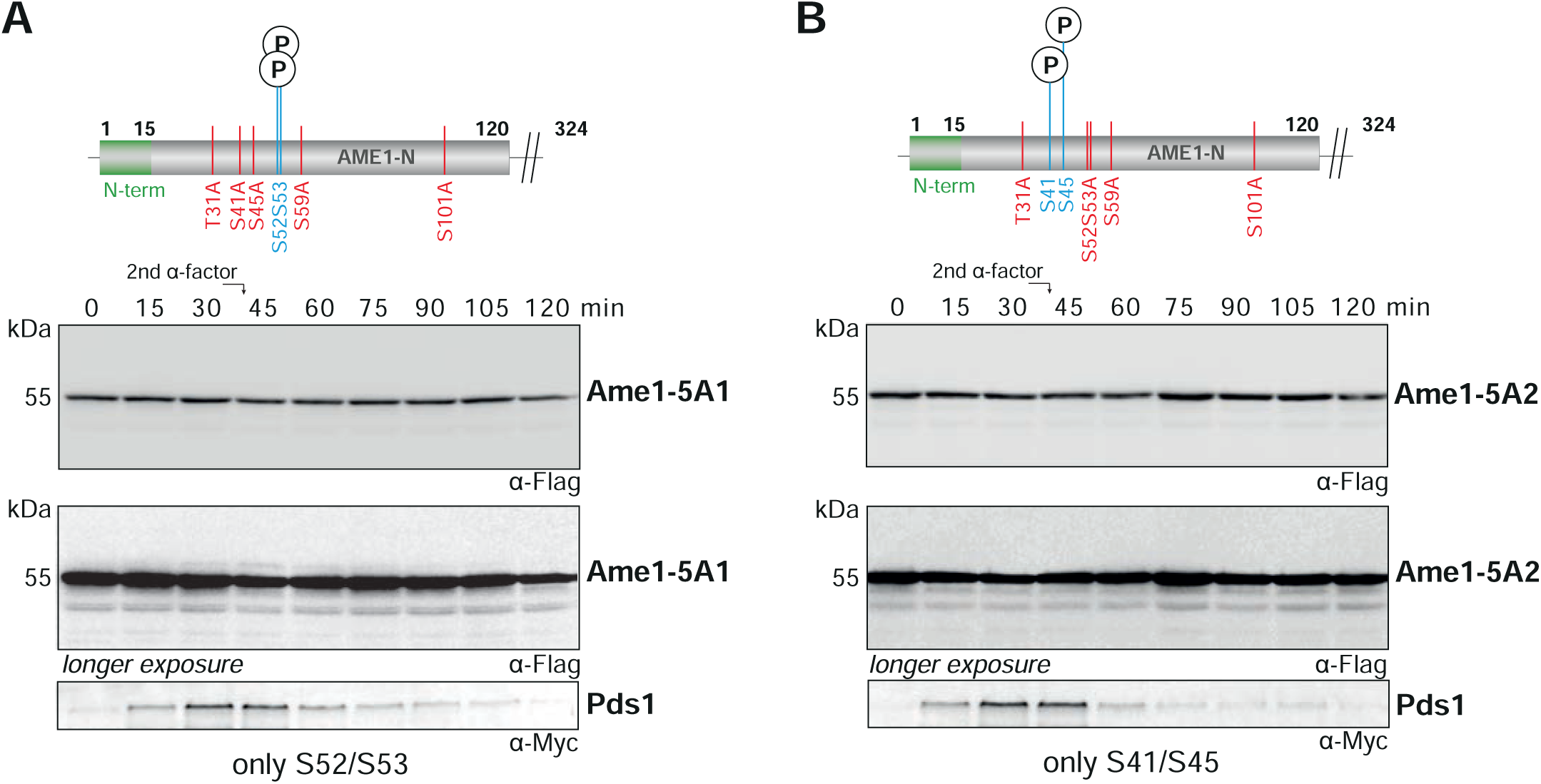
Additional analysis of endogenous Ame1 phospho-mutants over the cell cycle. **A:** A Flag-tagged Ame1 variant in which only phosphorylation of motif 1 was permitted (Ame1-5A1) was followed over the cell cycle by western blotting after release from alpha-factor arrest. **B:** A Flag-tagged Ame1 variant in which only phosphorylation of motif 2 was permitted (Ame1-5A2) was followed over the cell cycle by western blotting after release from alpha-factor arrest.

## Supplementary Movies

**Supplementary Movie 1**

Gaussian-accelerated Molecular Dynamics (GaMD) simulation of the doubly phosphorylated (S41 and S45) Ame1 peptide (red) binding to WD40 domain of Cdc4 (blue).

**Supplementary Movie 2**

GaMD simulation of the monophosphphorylated (S41) Ame1 peptide (red) binding to the WD40 domain of Cdc4 (blue).

**Supplementary Movie 3**

GaMD simulation of the monophosphphorylated (S45) Ame1 peptide (red) binding to the WD40 domain of Cdc4 (blue).

**Supplementary Movie 4**

GaMD simulation of the Ame1-CPD^ILTPP^ peptide (red), phosphorylated at T52, binding to the WD40 domain of Cdc4 (blue).

## References

Akiyoshi, B., Nelson, C.R., Duggan, N., Ceto, S., Ranish, J.A., and Biggins, S. (2013). The Mub1/Ubr2 ubiquitin ligase complex regulates the conserved Dsn1 kinetochore protein. PLoS Genet 9, e1003216.

Amato, A., Schillaci, T., Lentini, L., and Di Leonardo, A. (2009). CENPA overexpression promotes genome instability in pRb-depleted human cells. Mol Cancer 8, 119.

Anedchenko, E.A., Samel-Pommerencke, A., Tran Nguyen, T.M., Shahnejat-Bushehri, S., Popsel, J., Lauster, D., Herrmann, A., Rappsilber, J., Cuomo, A., Bonaldi, T., et al. (2019). The kinetochore module Okp1(CENP-Q)/Ame1(CENP-U) is a reader for N-terminal modifications on the centromeric histone Cse4(CENP-A). EMBO J 38.

Au, W.C., Zhang, T., Mishra, P.K., Eisenstatt, J.R., Walker, R.L., Ocampo, J., Dawson, A., Warren, J., Costanzo, M., Baryshnikova, A., et al. (2020). Skp, Cullin, F-box (SCF)-Met30 and SCF-Cdc4-Mediated Proteolysis of CENP-A Prevents Mislocalization of CENP-A for Chromosomal Stability in Budding Yeast. PLoS Genet 16, e1008597.

Biggins, S. (2013). The composition, functions, and regulation of the budding yeast kinetochore. Genetics 194, 817–846.

Bock, L.J., Pagliuca, C., Kobayashi, N., Grove, R.A., Oku, Y., Shrestha, K., Alfieri, C., Golfieri, C., Oldani, A., Dal Maschio, M., et al. (2012). Cnn1 inhibits the interactions between the KMN complexes of the yeast kinetochore. Nat Cell Biol 14, 614–624.

Case, D.A., Babin, V., Berryman, J.T., Betz, R.M., Cai, Q., Cerutti, D.S., Cheatham, T.E., III, Darden, T.A., Duke, R.E., Gohlke, H., et al. (2014). AMBER 14 (University of California, San Francisco).

Cheatham, T.E., III., Miller, J.L., Fox, T., Darden, T.A., and Kollman, P.A. (1995). Molecular Dynamics Simulations on Solvated Biomolecular Systems: The Particle Mesh Ewald Method Leads to Stable Trajectories of DNA, RNA and Proteins. J Am Chem Soc 117, 4193–4194.

Cheeseman, I.M., Chappie, J.S., Wilson-Kubalek, E.M., and Desai, A. (2006). The conserved KMN network constitutes the core microtubule-binding site of the kinetochore. Cell 127, 983–997.

Daniel, J.A., Yoo, J., Bettinger, B.T., Amberg, D.C., and Burke, D.J. (2006). Eliminating gene conversion improves high-throughput genetics in Saccharomyces cerevisiae. Genetics 172, 709–711.

Dhatchinamoorthy, K., Shivaraju, M., Lange, J.J., Rubinstein, B., Unruh, J.R., Slaughter, B.D., and Gerton, J.L. (2017). Structural plasticity of the living kinetochore. J Cell Biol 216, 3551–3570.

Dhatchinamoorthy, K., Unruh, J.R., Lange, J.J., Levy, M., Slaughter, B.D., and Gerton, J.L. (2019). The stoichiometry of the outer kinetochore is modulated by microtubule-proximal regulatory factors. J Cell Biol 218, 2124–2135.

Dimitrova, Y.N., Jenni, S., Valverde, R., Khin, Y., and Harrison, S.C. (2016). Structure of the MIND Complex Defines a Regulatory Focus for Yeast Kinetochore Assembly. Cell 167, 1014–1027 e1012.

Faustova, I., Bulatovic, L., Matiyevskaya, F., Valk, E., Ord, M., and Loog, M. (2021). A new linear cyclin docking motif that mediates exclusively S-phase CDK-specific signaling. EMBO J 40, e105839.

Feldman, R.M., Correll, C.C., Kaplan, K.B., and Deshaies, R.J. (1997). A complex of Cdc4p, Skp1p, and Cdc53p/cullin catalyzes ubiquitination of the phosphorylated CDK inhibitor Sic1p. Cell 91, 221–230.

Fischbock-Halwachs, J., Singh, S., Potocnjak, M., Hagemann, G., Solis-Mezarino, V., Woike, S., Ghodgaonkar-Steger, M., Weissmann, F., Gallego, L.D., Rojas, J., et al. (2019). The COMA complex interacts with Cse4 and positions Sli15/Ipl1 at the budding yeast inner kinetochore. Elife 8.

Gascoigne, K.E., Takeuchi, K., Suzuki, A., Hori, T., Fukagawa, T., and Cheeseman, I.M. (2011). Induced ectopic kinetochore assembly bypasses the requirement for CENP-A nucleosomes. Cell 145, 410–422.

Goh, P.Y., and Surana, U. (1999). Cdc4, a protein required for the onset of S phase, serves an essential function during G(2)/M transition in Saccharomyces cerevisiae. Mol Cell Biol 19, 5512–5522.

Haase, S.B., and Reed, S.I. (2002). Improved flow cytometric analysis of the budding yeast cell cycle. Cell Cycle 1, 132–136.

Hao, B., Oehlmann, S., Sowa, M.E., Harper, J.W., and Pavletich, N.P. (2007). Structure of a Fbw7-Skp1-cyclin E complex: multisite-phosphorylated substrate recognition by SCF ubiquitin ligases. Mol Cell 26, 131–143.

Haruki, H., Nishikawa, J., and Laemmli, U.K. (2008). The anchor-away technique: rapid, conditional establishment of yeast mutant phenotypes. Mol Cell 31, 925–932.

Heun, P., Erhardt, S., Blower, M.D., Weiss, S., Skora, A.D., and Karpen, G.H. (2006). Mislocalization of the Drosophila centromere-specific histone CID promotes formation of functional ectopic kinetochores. Dev Cell 10, 303–315.

Hinshaw, S.M., and Harrison, S.C. (2019). The structure of the Ctf19c/CCAN from budding yeast. Elife 8.

Holt, L.J., Tuch, B.B., Villen, J., Johnson, A.D., Gygi, S.P., and Morgan, D.O. (2009). Global analysis of Cdk1 substrate phosphorylation sites provides insights into evolution. Science 325, 1682–1686.

Homeyer, N., Horn, A.H.C., Lanig, H., and Sticht, H. (2006). AMBER force field parameters for phosphorylated amino acids in different protonation states: phosphoserine, phosphothreonine, phosphotyrosine and phosphohistidine. J Mol Model 12, 281–289.

Hornung, P., Troc, P., Malvezzi, F., Maier, M., Demianova, Z., Zimniak, T., Litos, G., Lampert, F., Schleiffer, A., Brunner, M., et al. (2014). A cooperative mechanism drives budding yeast kinetochore assembly downstream of CENP-A. J Cell Biol 206, 509–524.

Huis In ’t Veld, P.J., Jeganathan, S., Petrovic, A., Singh, P., John, J., Krenn, V., Weissmann, F., Bange, T., and Musacchio, A. (2016). Molecular basis of outer kinetochore assembly on CENP-T. Elife 5.

Jorgensen, W.L., Chandrasekhar, J., Madura, J.D., Impey, R.W., and Klein, M.L. (1983). Comparison of simple potential functions for simulating liquid water. J Chem Phys 79, 926–935.

Joung, I.S., and Cheatham, T.E., III. (2008). Determination of alkali and halide monovalent ion parameters for use in explicitly solvated biomolecular simulations. J Phys Chem B 112, 9020–9041.

Killinger, K., Bohm, M., Steinbach, P., Hagemann, G., Bluggel, M., Janen, K., Hohoff, S., Bayer, P., Herzog, F., and Westermann, S. (2020). Auto-inhibition of Mif2/CENP-C ensures centromere-dependent kinetochore assembly in budding yeast. EMBO J 39, e102938.

Klare, K., Weir, J.R., Basilico, F., Zimniak, T., Massimiliano, L., Ludwigs, N., Herzog, F., and Musacchio, A. (2015). CENP-C is a blueprint for constitutive centromere-associated network assembly within human kinetochores. J Cell Biol 210, 11–22.

Koivomagi, M., Valk, E., Venta, R., Iofik, A., Lepiku, M., Balog, E.R., Rubin, S.M., Morgan, D.O., and Loog, M. (2011). Cascades of multisite phosphorylation control Sic1 destruction at the onset of S phase. Nature 480, 128–131.

Kushnirov, V.V. (2000). Rapid and reliable protein extraction from yeast. Yeast 16, 857–860.

Lamiable, A., Thévenet, P., Rey, J., Vavrusa, M., Derreumaux, P., and Tufféry, P. (2016). PEP-FOLD3: faster de novo structure prediction for linear peptides in solution and in complex. Nucleic Acids Res 44, W449–W454.

Lampert, F., Mieck, C., Alushin, G.M., Nogales, E., and Westermann, S. (2013). Molecular requirements for the formation of a kinetochore-microtubule interface by Dam1 and Ndc80 complexes. J Cell Biol 200, 21–30.

London, N., Ceto, S., Ranish, J.A., and Biggins, S. (2012). Phosphoregulation of Spc105 by Mps1 and PP1 regulates Bub1 localization to kinetochores. Curr Biol 22, 900–906.

London, N., Raveh, B., Cohen, E., Fathi, G., and Schueler-Furman, O. (2011). Rosetta FlexPepDock web server - high resolution modeling of peptide-protein interactions. Nucleic Acids Res 39, W249–W253.

Lyons, N.A., Fonslow, B.R., Diedrich, J.K., Yates, J.R., 3rd, and Morgan, D.O. (2013). Sequential primed kinases create a damage-responsive phosphodegron on Eco1. Nat Struct Mol Biol 20, 194–201.

Lyons, N.A., and Morgan, D.O. (2011). Cdk1-dependent destruction of Eco1 prevents cohesion establishment after S phase. Mol Cell 42, 378–389.

Maier, J.A., Martinez, C., Kasavajhala, K., Wickstrom, L., Hauser, K.E., and Simmerling, C. (2015). ff14SB: improving the accuracy of protein side chain and backbone parameters from ff99SB. J Chem Theory Comput 11, 3696–3713.

Malvezzi, F., Litos, G., Schleiffer, A., Heuck, A., Mechtler, K., Clausen, T., and Westermann, S. (2013). A structural basis for kinetochore recruitment of the Ndc80 complex via two distinct centromere receptors. EMBO J 32, 409–423.

Maskell, D.P., Hu, X.W., and Singleton, M.R. (2010). Molecular architecture and assembly of the yeast kinetochore MIND complex. J Cell Biol 190, 823–834.

Miao, Y., Feher, V., and McCammon, J.A. (2015). Gaussian accelerated molecular dynamics: Unconstrained enhanced sampling and free energy calculation. J Chem Theor Comp 11, 3584–3595.

Musacchio, A., and Desai, A. (2017). A Molecular View of Kinetochore Assembly and Function. Biology (Basel) 6.

Nash, P., Tang, X., Orlicky, S., Chen, Q., Gertler, F.B., Mendenhall, M.D., Sicheri, F., Pawson, T., and Tyers, M. (2001). Multisite phosphorylation of a CDK inhibitor sets a threshold for the onset of DNA replication. Nature 414, 514–521.

Nishino, T., Rago, F., Hori, T., Tomii, K., Cheeseman, I.M., and Fukagawa, T. (2013). CENP-T provides a structural platform for outer kinetochore assembly. EMBO J 32, 424–436.

Ord, M., Moll, K., Agerova, A., Kivi, R., Faustova, I., Venta, R., Valk, E., and Loog, M. (2019a). Multisite phosphorylation code of CDK. Nat Struct Mol Biol 26, 649–658.

Ord, M., Venta, R., Moll, K., Valk, E., and Loog, M. (2019b). Cyclin-Specific Docking Mechanisms Reveal the Complexity of M-CDK Function in the Cell Cycle. Mol Cell 75, 76–89 e73.

Orlicky, S., Tang, X., Willems, A., Tyers, M., and Sicheri, F. (2003). Structural basis for phosphodependent substrate selection and orientation by the SCFCdc4 ubiquitin ligase. Cell 112, 243–256.

Ranjitkar, P., Press, M.O., Yi, X., Baker, R., MacCoss, M.J., and Biggins, S. (2010). An E3 ubiquitin ligase prevents ectopic localization of the centromeric histone H3 variant via the centromere targeting domain. Mol Cell 40, 455–464.

Roe, D.R., and Cheatham, T.E., III. (2013). PTRAJ and CPPTRAJ: Software for Processing and Analysis of Molecular Dynamics Trajectory Data. J Chem Theory Comput 9, 3084–3095.

Schleiffer, A., Maier, M., Litos, G., Lampert, F., Hornung, P., Mechtler, K., and Westermann, S. (2012). CENP-T proteins are conserved centromere receptors of the Ndc80 complex. Nat Cell Biol 14, 604–613.

Schwob, E., Bohm, T., Mendenhall, M.D., and Nasmyth, K. (1994). The B-type cyclin kinase inhibitor p40SIC1 controls the G1 to S transition in S. cerevisiae. Cell 79, 233–244.

Shao, J., Tanner, S.W., Thompson, N., and Cheatham, T.E., III. (2007). Clustering Molecular Dynamics Trajectories: 1. Characterizing the Performance of Different Clustering Algorithms. J Chem Theory Comput 3, 2312–2334.

Singh, P., Pesenti, M.E., Maffini, S., Carmignani, S., Hedtfeld, M., Petrovic, A., Srinivasamani, A., Bange, T., and Musacchio, A. (2021). BUB1 and CENP-U, Primed by CDK1, Are the Main PLK1 Kinetochore Receptors in Mitosis. Mol Cell 81, 67–87 e69.

Skowyra, D., Craig, K.L., Tyers, M., Elledge, S.J., and Harper, J.W. (1997). F-box proteins are receptors that recruit phosphorylated substrates to the SCF ubiquitin-ligase complex. Cell 91, 209–219.

Swaney, D.L., Beltrao, P., Starita, L., Guo, A., Rush, J., Fields, S., Krogan, N.J., and Villen, J. (2013). Global analysis of phosphorylation and ubiquitylation cross-talk in protein degradation. Nat Methods 10, 676–682.

Tanaka, K., Kitamura, E., Kitamura, Y., and Tanaka, T.U. (2007). Molecular mechanisms of microtubule-dependent kinetochore transport toward spindle poles. J Cell Biol 178, 269–281.

Tien, J.F., Umbreit, N.T., Gestaut, D.R., Franck, A.D., Cooper, J., Wordeman, L., Gonen, T., Asbury, C.L., and Davis, T.N. (2010). Cooperation of the Dam1 and Ndc80 kinetochore complexes enhances microtubule coupling and is regulated by aurora B. J Cell Biol 189, 713–723.

Van Hooser, A.A., Ouspenski, II, Gregson, H.C., Starr, D.A., Yen, T.J., Goldberg, M.L., Yokomori, K., Earnshaw, W.C., Sullivan, K.F., and Brinkley, B.R. (2001). Specification of kinetochore-forming chromatin by the histone H3 variant CENP-A. J Cell Sci 114, 3529–3542.

Wallace, A.C., Laskowski, R.A., and Thornton, J.M. (1995). LIGPLOT: a program to generate schematic diagrams of protein-ligand interactions. Protein engineering, design and selection 8, 127–134.

Yan, K., Yang, J., Zhang, Z., McLaughlin, S.H., Chang, L., Fasci, D., Ehrenhofer-Murray, A.E., Heck, A.J.R., and Barford, D. (2019). Structure of the inner kinetochore CCAN complex assembled onto a centromeric nucleosome. Nature 574, 278–282.

